# Mycobacterium tuberculosis methyltransferase perturbs host epigenetic programming to promote bacterial survival

**DOI:** 10.1101/2023.02.24.529973

**Authors:** Prakruti R Singh, Venkatareddy Dadireddy, Shubha Udupa, Shashwath Malli Kalladi, Somnath Shee, Sanjeev Khosla, Raju S Rajmani, Amit Singh, Suryanarayanarao Ramakumar, Valakunja Nagaraja

## Abstract

*Mycobacterium tuberculosis* (*Mtb*) has evolved several mechanisms to counter host defense arsenal for its proliferation. We show that *Mtb* employs multi-pronged approach to modify host epigenetic machinery for its survival upon infection. It secretes methyltransferase (MTase) Rv2067c into macrophages, trimethylating K79 of histone H3 in non-nucleosomal context. Rv2067c downregulates host MTase DOT1L, decreasing its H3K79me3 mark added nucleosomally on pro-inflammatory response genes. Consequent inhibition of caspase8 dependent apoptosis and enhancement of RIPK3 mediated necrosis results in increased pathogenesis. In parallel, Rv2067c enhances the expression of SESTRIN3, NLRC3 and TMTC1 enabling the pathogen to overcome host inflammatory and oxidative response. We provide structural basis for differential methylation of H3 by Rv2067c and DOT1L. The structure of Rv2067c and DOT1L explain how their action on H3K79 is temporally and spatially separated enabling Rv2067c to effectively intercept the host epigenetic circuit and downstream signalling.

## Introduction

Pathogen infection leads to activation of several host defence mechanisms such as immune response, oxidative burst, inflammation, cell death/survival to curtail the infection^1^. This host pathogen interaction result in alteration of transcriptomic landscape of the infected cell leading to selective activation and repression of genes, influencing different signalling pathways^2^. Amongst many, a key mechanism emerging is the crosstalk between epigenetic modifications and the transcription machinery^1, 3, 4^.

*Mycobacterium tuberculosis (Mtb),* a successful intracellular pathogen and a leading cause of mortality^5^, has evolved diverse mechanisms in response to multiple stresses encountered upon infection. It inhibits oxidative and nitrosative stress, supresses apoptosis and immune response, also manipulating host ubiquitin system for its enhanced survival^6–9^. About 5% of *Mtb* genome encodes epigenetic modifiers that include kinases, methyltransferases, acetyltransferases and succinyltransferases hinting their role in manipulation of epigenetic signalling post infection (p.i). A few studies indicate the role of these modifiers in fine-tuning the mycobacterial epigenome^10–14^. However, a majority of them remain uncharacterized and their involvement in modifying bacterial or host epigenome is yet to be understood.

Here, we have investigated the structure and function of a *Mtb* methyltransferase (MTase) Rv2067c. Our study revealed that Rv2067c is secreted by *Mtb* into the macrophages where it exerts its function by at least two mechanisms. First, it trimethylates free histone H3 at lysine 79 (H3K79) to alter the expression of a gene subset in the host. Second, it represses host methyltransferase DOT1L expression pre-empting DOT1L activity on H3K79 in nucleosomal context. We provide structural basis on how Rv2067c methylates free histone H3 and not nucleosomal H3, by comparing its structure with that of nucleosome bound DOT1L. This unprecedented multi-pronged epigenetic strategy elicited by *Mtb* through its effector leads to simultaneous increase in necrosis and inhibition of apoptosis ensuring pathogen’s survival.

## Results

### Rv2067c methylates Histone H3

Rv2067c was identified as one of the interacting partners of *Mtb* histone like protein HU (MtHU) in pulldown experiments with MtHU as a bait (Extended Data Fig. 1a). Sequence analysis of Rv2067c revealed the presence of SAM dependent MTase domain suggesting Rv2067c is a MTase (Extended Data Fig. 1b). However, on performing MTase assays, Rv2067c did not methylate recombinant MtHU (Extended Data Fig. 1c). As MtHU carboxy-terminal domain (CTD) shows remarkable similarity with histone tails^15^, we investigated whether mammalian histones are targets for the MTase activity of Rv2067c. In MTase assays, using salt extracted histones from THP1 monocytes as substrates, Rv2067c as MTase and tritiated SAM as methyl donor, methylation of histones was detected (Fig. 1a and Extended Data Fig. 1d). To determine which of the histones is a substrate for Rv2067c, MTase assays with individual recombinant mammalian histones was performed. Among the four histones, only recombinant H3 was methylated by Rv2067c (Fig. 1b and c). Using tandem MS/MS, the site of methylation was identified as lysine 79 of H3 (H3K79). A mass shift of 42Da at lysine 79 of H3 implied Rv2067c trimethylates H3K79 (Fig. 1d). To ascertain, MTase assays were carried out with unmodified 73-83aa peptide of H3, followed by dot blot. H3K79me3 antibody detected a signal, confirming Rv2067c trimethylates H3K79 (Fig. 1e). Further, when lysine 79 of H3 was mutated to alanine, H3A79 protein was not methylated by Rv2067c (Fig. 1f and Extended Data Fig. 1e), re-confirming its specificity for H3K79. To demonstrate SAM dependent activity of Rv2067c, conserved GxG motif was mutated to RxR and MTase assay showed that the mutation abolished the activity of Rv2067c-RxR (Fig 1g and Extended data Fig. 1f) Intriguingly, H3K79 methylation is a conserved epigenetic mark in eukaryotes catalysed by DOT1L. Yeast Dot1 and hDOT1L catalyze mono-, di- and tri-methylation of H3K79, but only in the nucleosome core particle (NCP) and free H3 is not the substrate^16, 17^. To determine whether Rv2067c can also methylate nucleosomal H3, we performed MTase reactions with reconstituted H3-H4 dimer, histone octamers and NCPs (Extended Data Fig. 1g and h). Rv2067c methylated only free H3 and did not methylate H3 in octamer or nucleosomal core. (Fig. 1h, Extended Data Fig. 1i).

**Fig. 1:**
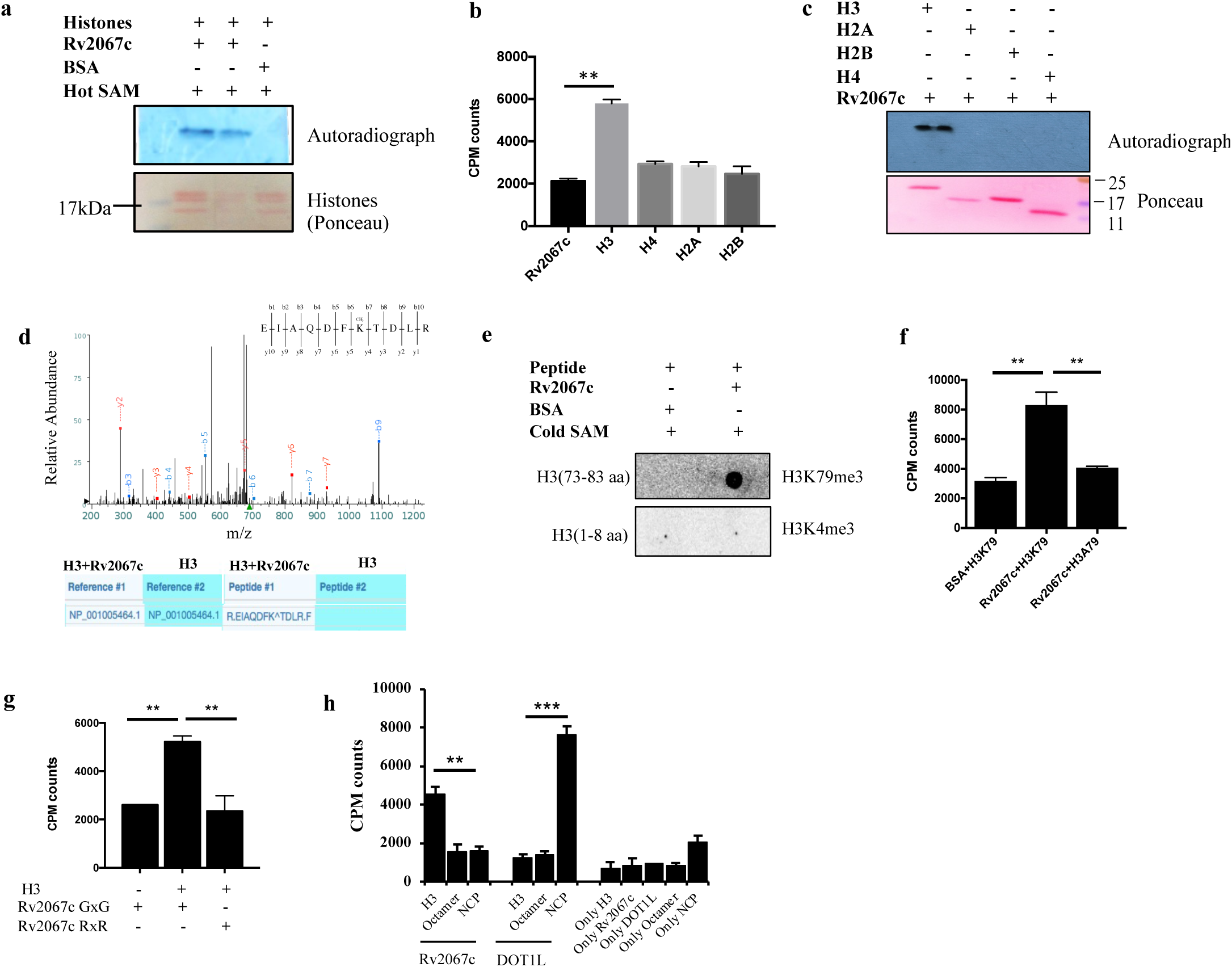
Rv2067c *in vitro* methylates histone H3 at lysine 79. **a,** Autoradiograph shows methylation activity of Rv2067c with histones salt extracted from THP1 monocytes as substrate and tritiated SAM as a methyl group donor (Lane 1 and 2). BSA (lane 3) was kept as negative control. Ponceau staining of blot shows loading of histones. **b,** Bar graph depicts scintillation counts (CPM) for MTase assays with individual recombinant mammalian histones as substrate, Rv2067c as MTase and tritiated SAM as a methyl group donor. Recombinant Rv2067c was kept as negative control. **c,** Autoradiograph shows methylation activity of Rv2067c against recombinant H3, H2A, H2B and H4 as substrate and tritiated SAM as a methyl group donor. Ponceau staining of blot shows loading of histones. **d,** MS/MS spectra of H3K79 trimethylation performed by Rv2067c on recombinant H3. The table below the spectra shows peptide EIAQDFK^TDLR was trimethylated (^ represents trimethylation) when assay was performed with Rv2067c and H3. Only H3 did not show lysine 79 trimethylation. **e,** Blots depicts *in vitro* MTase assays with unmodified peptide corresponding to H3 73-83 aa and 1-8 aa as substrate, Rv2067c as MTase and cold SAM as a methyl group donor. Blot was probed with H3K79me3 and H3K4me3 antibodies respectively. Peptide corresponding to H3 1-8 aa was kept as control. **f,** Scintillation counts for *in vitro* MTase assays with recombinant H3 and H3A79 mutant protein as substrate and Rv2067c as MTase. BSA was kept as negative control. **g,** Scintillation counts for *in vitro* MTase assays with recombinant H3 as substrate and Rv2067c GxG and Rv2067c RxR mutant protein as MTase. **h,** *In vitro* MTase assay with Rv2067c as MTase and recombinant H3, reconstituted octamers and NCPs as substrate. Recombinant DOT1L was kept as positive control for MTase assay with NCPs. Recombinant H3, Rv2067c, DOT1L, octamers and NCPs were kept as negative control. The data for MTase assay (scintillation counts) is representative of three independent experiments and plotted as mean. Error bars correspond to s.d and **,P-value <0.01, ***,P-value <0.001(Student’s t-test).

### Structure of Rv2067c differs from that of DOT1L

To understand the structural differences conferring the context-dependent methylation of H3K79 by DOT1L and Rv2067c and the basis of free H3 methylation by Rv2067c, we determined the structure of Rv2067c to 2.40 Å resolution (Extended Data Table 1). Each monomer (407aa) of Rv2067c homodimer (Fig. 2a, Extended Data Fig. 2a) is composed of three domains: N-terminal SAM-binding catalytic domain (CD), dimerization domain (DD), and C-terminal domain (CTD) (Fig. 2b, c, d). The catalytic domain contains a central seven-stranded β-sheet flanked by α-helices, a characteristic of seven-β-strand (7BS) MTases (class-I MTases)^18^ (Fig. 2c and Extended Data Fig. 2c). The DD is composed of two subdomains, separated on the primary sequence, namely large subdomain (LSD; residues 163-230, four α-helices) and small subdomain (SSD; residues 257-291, two α-helices), and are insertions in the catalytic domain at β5-α5 and β6-β7 loops, respectively (Fig. 2c, d and Extended Data Fig. 2c). Dimer is formed by the cross-subdomain interactions between the DDs of the two monomers A and B in a fashion: LSD_A_-SSD_B_ and LSD_B_-SSD_A_, and LSD_A_-LSD_B_ (Fig. 2b). CTD is about 100 residues long with α + β architecture (Fig. 2c, d and Extended Data Fig. 2c). The dimerization interface in conjunction with other two domains forms two parallel trough-like structures, across the dimer, the putative acceptor substrate-binding sites (Fig. 2a, b and Fig. 3a).

**Fig. 2:**
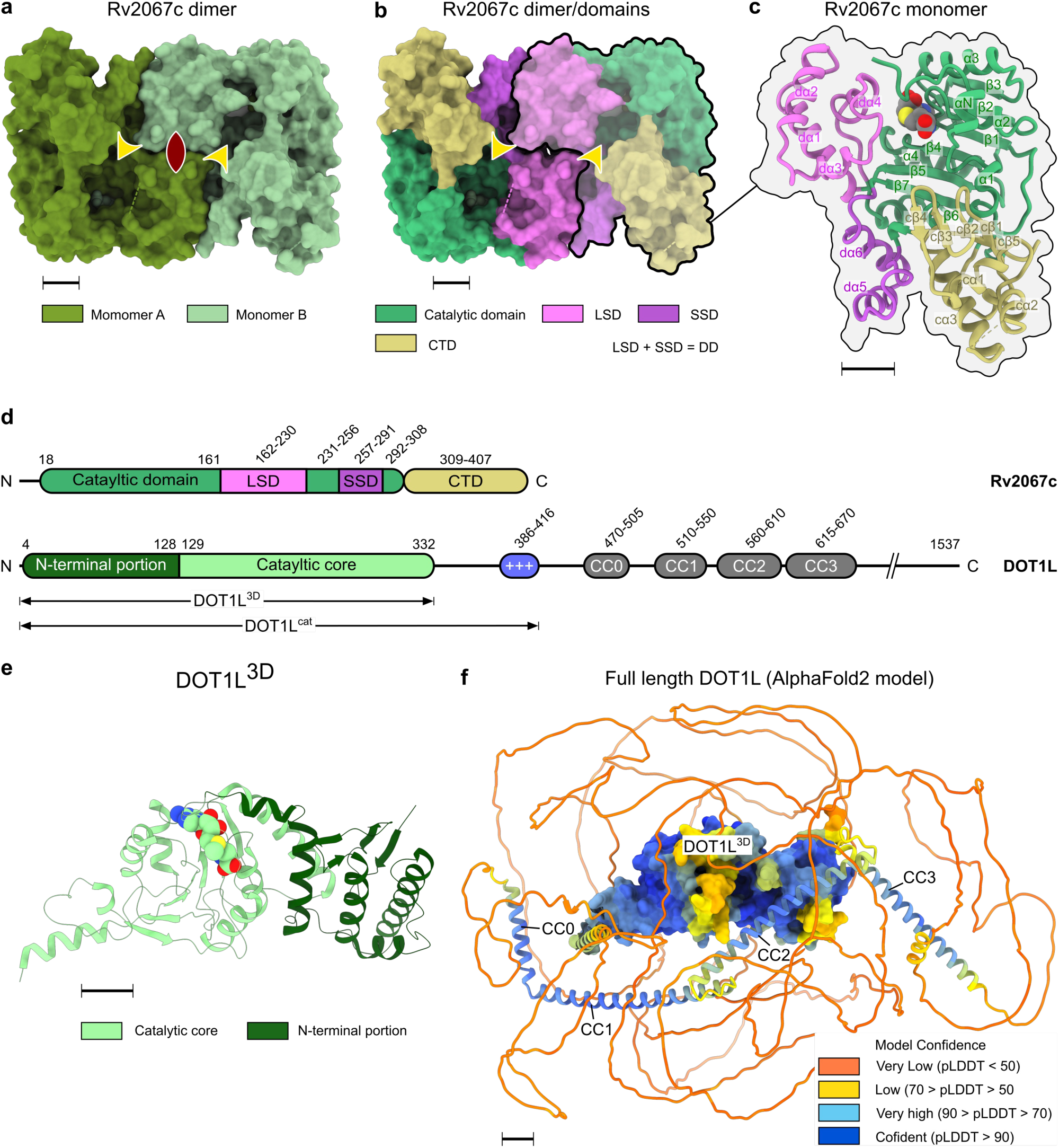
Structure of Rv2067c and its comparison to hDOT1L. **a,** Crystal structure of Rv2067c. The asymmetric unit contains two molecules of Rv2067c related by two-fold symmetry (homodimer). The two-fold axis is perpendicular to the plane and shown as biconvex symbol. **b** and **c,** Each monomer of Rv2067c contains a catalytic domain, dimerization domain (DD, composed of large subdomain, LSD and small subdomain, SSD) and C-terminal domain (CTD). Dimer is formed by cross subdomainal interactions between the DDs. SAH is rendered as spheres. (a and b) the dashed lines represent the missing electron density (disordered regions). The putative substrate-binding troughs are indicated with yellow arrow heads. **d,** Schematic of domain organization in Rv2067c (top) and DOT1L (bottom). e, Crystal structure of DOT1L (PDB: 1NW3). Residues 5-332 are structured (DOT1L3D). The DOT1L3D contains an N-terminal portionand catalyticcore. SAM is rendered as spheres. f, Full-length human DOT1L (1537 aa) structure model (generated using AlphaFold2). The structured part of the catalytic domain (DOT1L3D) is shown as surface and the remaining region as cartoon which is disordered with interspersed α-helices that form coiled-coil structures with the interacting partners. The model is colored according to the AlphaFold2 prediction confidence. The low confidence score indicates intrinsic disorder. Scalebars 10 Å.

**Fig. 3:**
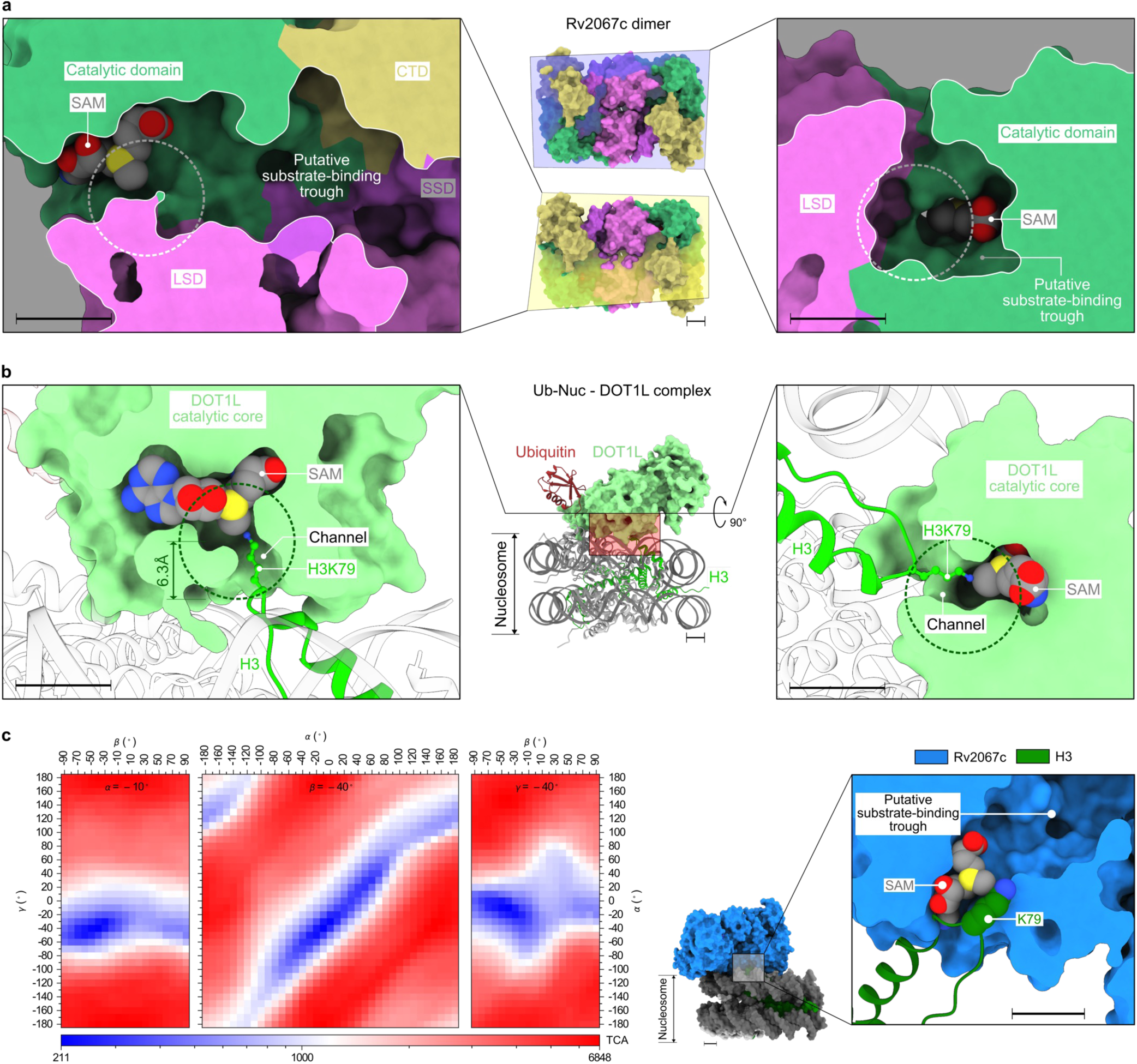
Comparison of substrate binding regions between Rv2067c and Dot1L. **a,** Cross sections (left and right panels) of putative substrate-binding trough of Rv2067c as depicted by planes passing through Rv2067c dimer (middle panel). The active site lies deep inside (right panel) at one end (left panel) of the trough and encompassed by catalytic domain and LSD (right panel). The region opposite to methyl group of SAM is occluded, that leaves no room for substrate lysine binding. Rv2067c is surface rendered. **b,** The complex between ubiquitinated nucleosome and DOT1L^cat^ (middle panel). The substrate lysine (H3K79 of nucleosome) binds in a narrow channel (cross sections across SAM, left and right panels). The length of the channel is about the length of the lysine sidechain in its extended rotamer. Nucleosome and ubiquitin are depicted as cartoons, Dot1L is surface rendered. The substrate lysine outreach to the SAM methyl group, in both a and b, are encircled. **c,** The total number of clashing atoms (TCA) for each pose are shown as a function of elemental rotation angles (α, β or γ) (left panel). Each subplot corresponds to the value of either α or β or γ at minimum TCA value (i.e., 211). The pose between Rv2067c and nucleosome reaction complex with minimum TCA (right panel). In this pose nucleosomal H3K79 accesses the reaction-center of Rv2067c only via the SAM entry site.

Structures of DOT1L in solo and in complex with nucleosome provide the structural basis for nucleosomal methylation by DOT1L^19–23^. In contrast to dimeric Rv2067c, DOT1L is 1,537 aa long monomer with catalytically active region spanning the first 416 residues (DOT1L^cat^), of which residues 4-332 are structured (DOT1L^3D^). DOT1L^3D^ is an elongated structure with an N-terminal portion and a SAM-binding catalytic core with a 7BS MTase fold (Fig. 2d, e, f and Extended Data Fig. 2c). DOT1L^3D^ is followed by a positively charged lysine-rich region (residues 386-416) involved in DNA binding and a disordered region with interspersed coiled-coil domains that aid in protein-protein interactions^19, 20, 24, 25^ (Fig. 2d, f). Barring a common class-I MTase fold (Extended Data Fig. 2d), Rv2067c and DOT1L markedly differ in their domain composition, architecture, and oligomeric state.

### Rv2067c structure precludes nucleosomal H3K79 methylation

Apart from the insights obtained from the gross structural differences, mechanism of nucleosomal methylation by DOT1L and comparison of active sites of DOT1L and Rv2067c provided the basis for substrate-context dependent methylation by Rv2067c. During methylation, DOT1L binds across the nucleosome (Nuc) or ubiquitinated nucleosome (UbNuc) and interacts with different regions of the Nuc or UbNuc. These include (1) hydrophobic interaction between the C-terminal helix of DOT1L^3D^ and ubiquitin, (2) contact between DOT1L R282 and H2A-H2B acidic patch, (3) tucking of basic residues of H4 tail into the acidic groove of DOT1L N-terminal portion, and (4) interaction between H3K79 loop, and F131 and W305 loops of DOT1L (Fig. S1a). These structural elements of DOT1L that aid in nucleosomal methylation and their equivalences are different in Rv2067c with respect to the primary sequence and tertiary structure (Extended Data Fig. 3). In addition, DOT1L binding stimulates a conformational transition in H3K79 loop which places H3K79 in the active site of DOT1L from its inaccessible conformation, which is pivotal for methylation ^20–23^ (Fig. S1a).

Notably, the active site architectures of Rv2067c and DOT1L are starkly different. The active-site of DOT1L is a narrow channel, formed by three loops viz. a loop connecting N-terminal portion and catalytic core (F131 loop), β4-α4 and β6-β7 (W305 loop), enough to accommodate H3K79 side chain. At one end of the channel lies the methyl moiety of SAM and the other end is open for H3K79 entry^19, 20^ (Fig. S1b and Fig. 3b). In contrast, Rv2067c contains a putative substrate-binding trough formed by all three domains (Fig. 2b and Fig. 3a). The active site lies at one end of the trough, deep inside the protein, encompassed by LSD and CD (Fig. 3a). Such an active site architecture presumably obstructs the accessibility by bulky regions of the substrate (nucleosomal H3K79) but could allow extended substrates like free H3. By a computational procedure (rotation scan) using enzyme-substrate reaction-complex model (Methods), the accessibility of Rv2067c active site by nucleosome was assessed with a supposition that the active-state H3K79 conformation exists in nucleosomes. The rotation-scan calculated a minimal number of 211 atoms involved in atomic clashes in Rv2067c-nucleosome reaction-complex and in the corresponding pose, nucleosome approaches Rv2067c active site via an unconventional direction, the SAM entry site, instead of putative substrate-binding site (Fig. 3c). In short, due to the lack of nucleosome interacting structural elements and different active site architectures, Rv2067c is incompetent to methylate nucleosomal H3K79 but its putative substrate-binding trough possibly accommodates and methylates free H3.

### Active site dynamics facilitate free H3 methylation (requires suggestions)

The substrate-binding trough of Rv2067c resembles the substrate-binding cleft of protein lysine methyltransferase 1 and 2 (PKMT1/2) from rickettsial species^26^ (Supplementary information). Inspection of Rv2067c substrate-binding trough revealed that the residues facing the methyl moiety of SAM occlude the H3K79 binding (Fig. 3a and Fig. S2). Sequence analysis showed that these residues are conserved (Extended Data Fig. 4, Supplementary information) and constitute a putative active site (H3K79 binding site). Structural analysis showed that the active site of protein MTases lies nearly perpendicular to the SAM binding pocket and accommodates the substrate residue (Fig. S2). The relatively high temperature factors (>90 Å^2^) of a moderately (Q228) and highly conserved (Y20) residues indicated the active site is dynamic in nature. During a 100 ns long all-atom molecular dynamics (MD) simulations, opening of the occluded active site pocket was observed that makes room for H3K79 (Fig. 4, Supplementary Movie). In addition to the active site dynamics, the disordered dα3-dα4 loop (residues 209-214, part of LSD and the substrate-binding trough) (Fig. 2a, b and Fig. S3) is likely to also facilitate H3 binding to the substrate-binding trough by an induced fit.

**Fig. 4:**
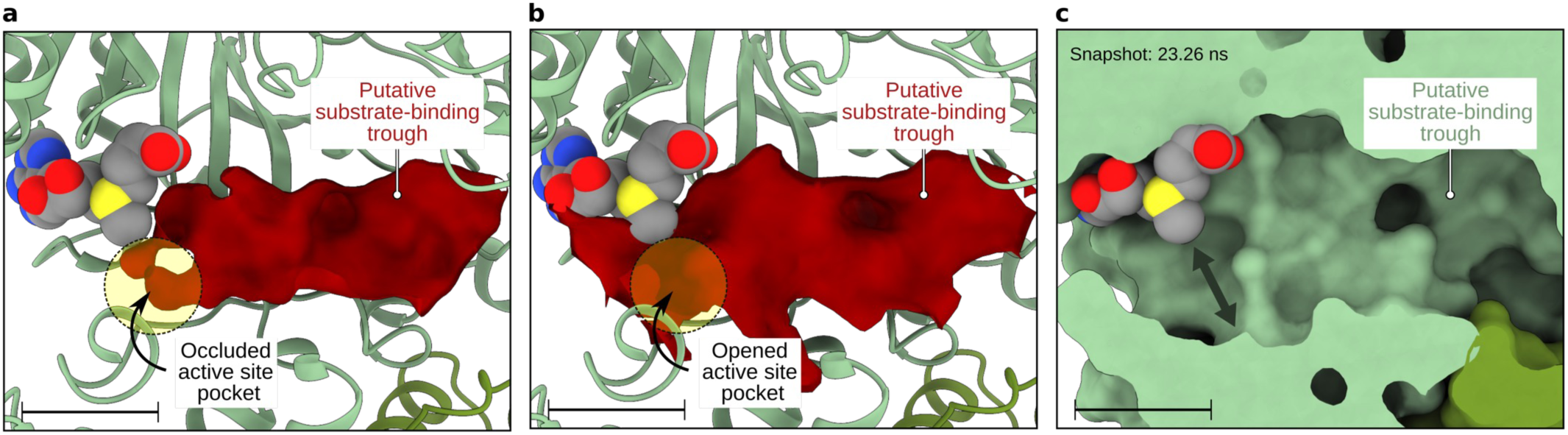
Rv2067c active site dynamics. Volumetric density map of Rv2067c putative substrate-binding trough calculated for a crystal structure and MD trajectory. **a,** In the crystal structure, the active site is occluded. **b,** During 100 ns MD simulations, the active site pocket is opened to accommodate substrate lysine. The volumetric density map is shown as an isosurface at 0.1 contour level. **c,** The opened active site (marked with double headed arrow) for a simulation snap short is shown. Scalebars 10 Å.

For modeling Rv2067-H3 complex, we resorted to a 11-mer H3 peptide 73-EIAQDFKTDLR-83 due to the lack of free H3 tertiary structure information (Supplementary information). This peptide is a competent substrate for Rv2067c (Fig. 1e) and assumes a random coil structure while part of a free H3^27, 28^ (Fig. S4a). PKMT1/2 also methylate multiple lysine residues in the random coil regions of the Outer membrane protein B (OmpB)^29^. It is also clear that the globular structures face steric hindrance to access the deep active sites via narrow trough. Hence, we modeled H3 peptide into the substrate-binding trough of Rv2067c based on the active site dynamics (Fig. 4) and the mode of substrate binding to MTases (Fig. S2). Due to the directional ambiguity, as the peptide binds in its extended conformation in a narrow trough, we propose two modes of peptide binding, i.e., N- to C-terminal or C- to N-terminal direction with respect to the active site (Fig. 5). The binding conformations resemble the binding of histone tails to SET domain MTases^30, 31^ or protein arginine N-MTases (PRMTs)^32^. In both the modes, side chain of H3K79 occupied the active site and oriented near perpendicular to the SAM axis as seen in protein MTase complexes (Fig. 5), amenable for methylation.

**Fig. 5:**
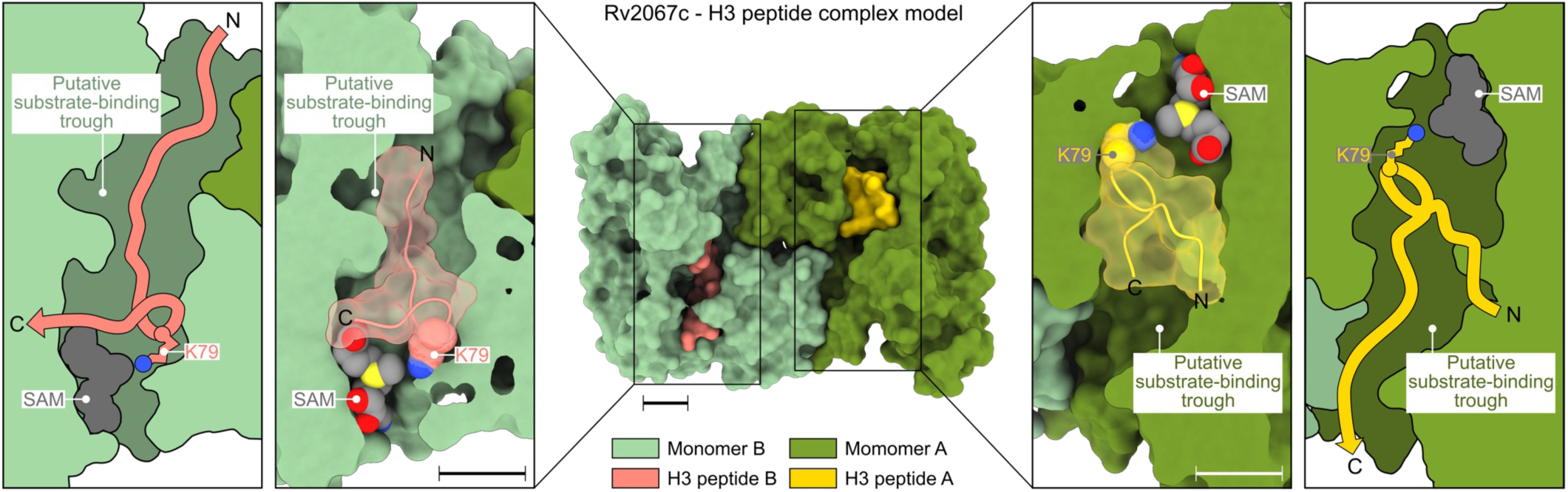
Rv2067c-H3 peptide complex model. Rv2067c-H3 peptide complex model (middle panel) depicting H3 peptide binding in the putative substrate-binding trough. The H3 peptide was modeled in two binding modes, along the trough, N- to C-terminal (left panel) or C- to N-terminal (right panel) with respect to the active site. In both modes, H3K79 can access the SAM methyl group in a similar fashion seen in MTase-substrate complexes (Fig. S2). Scalebars 10 Å.

### Rv2067c trimethylates H3K79 upon *Mtb* infection

Having established H3 as a substrate of Rv2067c, we investigated the action of Rv2067c upon *Mtb* infection in human macrophages. First, we checked the interaction of Rv2067c with endogenous H3 by transfecting HEK293T with pcDNA:Rv2067c-SFB (S-protein,FLAG,streptavidin-binding peptide). Immunoblot showed H3 interacts with Rv2067c upon pulldown (Fig. 6a). Interaction between Rv2067c and H3 was re-confirmed by performing IP with H3 antibody (Extended Data Fig. 5a). Enhanced H3K79me3 in cells expressing Rv2067c (Fig. 6b) compared to Rv2067cRxR indicated methylation by the MTase (Fig. 6c). To confirm methylation of cytoplasmic H3 by Rv2067c, HEK293T were co-transfected with FLAG tagged H3 (HEK:H3-FL) and Rv2067c. When nuclear and cytoplasmic extracts were analysed, H3K79me3 was observed on cytosolic H3, while DOT1L specific H3K79me3 was observed in the nuclear fraction (Fig. 6d).

**Fig. 6:**
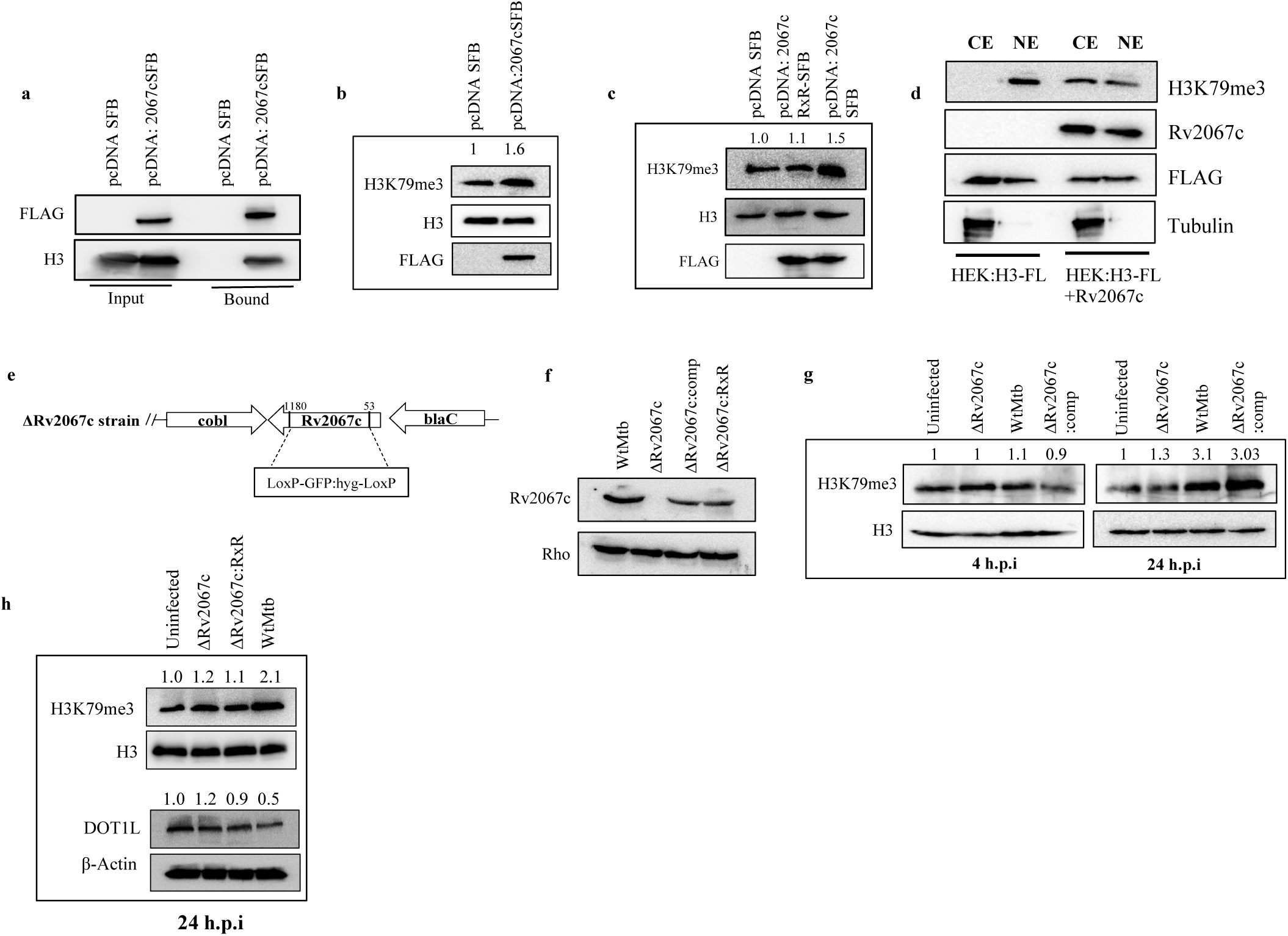
Rv2067c methylates histone H3 at lysine 79 upon infection in THP1 macrophages. **a,** Western blot depicts interaction between Rv2067c and H3. Pull down was performed with streptavidin beads on lysates of HEK293T transfected with pcDNA: Rv2067cSFB or pcDNA SFB (control) constructs. 5% of whole cell lysate was kept as Input. Blots were probed with FLAG and H3 antibodies. **b,** Immunoblot shows H3K79me3 mark in cell lysate of HEK293T transfected with pcDNA: Rv2067cSFB and pcDNA SFB construct. Blot was probed with H3K79me3 antibody. H3 was used as loading control and FLAG shows expression of Rv2067c. **c,** Immunoblot shows H3K79me3 mark for the cell lysate of HEK293T transfected with pcDNA SFB (lane1), pcDNA: Rv2067cRxR-SFB (lane2) and pcDNA: Rv2067cSFB (lane3). Blot was probed with H3K79me3 antibody. H3 was used as loading control. FLAG depicts expression of Rv2067c. **d,** Western blot depicts H3K79me3 mark added by Rv2067c in the cytosolic fraction of HEK293T expressing Rv2067c and FLAG tagged H3 (H3-FL). HEK293T cells expressing HEK:H3-FL were kept as control. Blot was probed with FLAG and Rv2067c antibodies to show expression of H3 and Rv2067c respectively. Tubulin was used as control for cytosolic fraction. NE-nuclear extract, CE-cytosolic extract. **e,** Schematic representation of the strategy used for the generation of *Rv2067c* gene knockout. The internal fragment of Rv2067c from +53 bp to +1180 bp was deleted and replaced with hygromycin resistance cassette (LoxP-GFP:hyg-LoxP). **f,** Western blot depicts Rv2067c levels in the cell lysates of Wt*Mtb* (lane1), ΔRv2067c (lane 2), ΔRv2067c:comp (lane 3) and ΔRv2067c:RxR (lane 4). Blot was probed with Rv2067c and Rho (control) antibodies. **g,** Western blot depicts H3K79me3 mark in THP1 cell lysates post 4 and 24 hours of infection with various *Mtb* strains. H3 was used as loading control. Lane1: uninfected THP1 macrophages; Lane 2, 3 and 4: THP1 macrophages infected with ΔRv2067c, Wt*Mtb* And ΔRv2067c:comp strains respectively. **h,** Immunoblot depicts level of H3K79me3 mark and DOT1L in THP1 cell lysates 24 h.p.i with ΔRv2067c:RxR. H3 and β-Actin were used as loading control respectively. Lane1: uninfected THP1 macrophages; Lane 2, 3 and 4: THP1 macrophages infected with ΔRv2067c, ΔRv2067c:RxR and Wt*Mtb* strains respectively. **f-h**, Comp stands for complemented Values above the blot represent quantitation (arbitary units).

To probe Rv2067c function upon *Mtb* infection, knockout (ΔRv2067c), complemented (ΔRv2067c:comp) and over-expression strains (Rv2067c:OE) were constructed. Schematic depicts knockout generation strategy (Fig. 6e). Knockout and complemented strains were confirmed by PCR, qRT-PCR (Extended Data Fig. 5b-e) and immuno-blotting with Rv2067c antibody (Fig. 6f). These strains did not show growth difference *in vitro* (Extended Data Fig. 5g). When macrophages were infected with these strains, enhanced H3K79me3 mark was observed for Wt*Mtb* and ΔRv2067c:comp, with no change in the knockout (Fig. 6g) and ΔRv2067c:RxR strain (Fig. 6h), 24hr post infection (h.p.i).Thus, it is apparent that Rv2067c, a SAM dependent MTase, not only tri-methylates H3K79 *in vitro* but also upon transfection and *Mtb* infection in macrophages.

### Rv2067c is secreted into the host macrophages

Since Rv2067c trimethylates H3K79 in the host, we examined its secretion from mycobacteria. Rv2067c is secreted into the culture filtrate of Wt*Mtb* and *M.smeg*:Rv2067c-FLAG (Extended Data Fig. 6a and b). As *M.smeg* does not have a Rv2067c homolog, its secretory nature was validated by detecting peptides specific to Rv2067c in the culture filtrate of *M.smeg*:Rv2067c-FLAG by mass spectrometry analysis (Extended Data Fig. 6d). Confocal microscopy p.i revealed MTase’s localization both in the cytoplasm and nucleus of macrophages infected with Rv2067c:OE and *M.smeg*:Rv2067c-FLAG (Extended Data Fig. 6e and f). Subcellular fractionation of HEK293T transfected with Rv2067c confirmed its localization in cytoplasm and nucleus (Extended Data Fig. 6g).

### Rv2067c modulates expression of DOT1L

The increase in H3K79me3 upon infection is likely due to the activity of Rv2067c. Alternately the increase could be DOT1L specific as both MTases target the same site in H3. Hence, we examined the expression of DOT1L upon *Mtb* infection. Surprisingly, DOT1L expression was reduced in macrophages infected with Wt*Mtb* and Rv2067c:comp, 12 and 24 h.p.i (Fig. 7a and b and Extended Data Fig. 7b) and also in HEK293T transiently expressing Rv2067c (Extended Data Fig. 7a). However, DOT1L expression did not alter in macrophages infected with ΔRv2067c (Fig. 7a and b) and ΔRv2067c:RxR, 12 and 24 h.p.i (Fig. 6h). These results imply that the higher level of H3K79me3 is likely to be Rv2067specific.

**Fig. 7:**
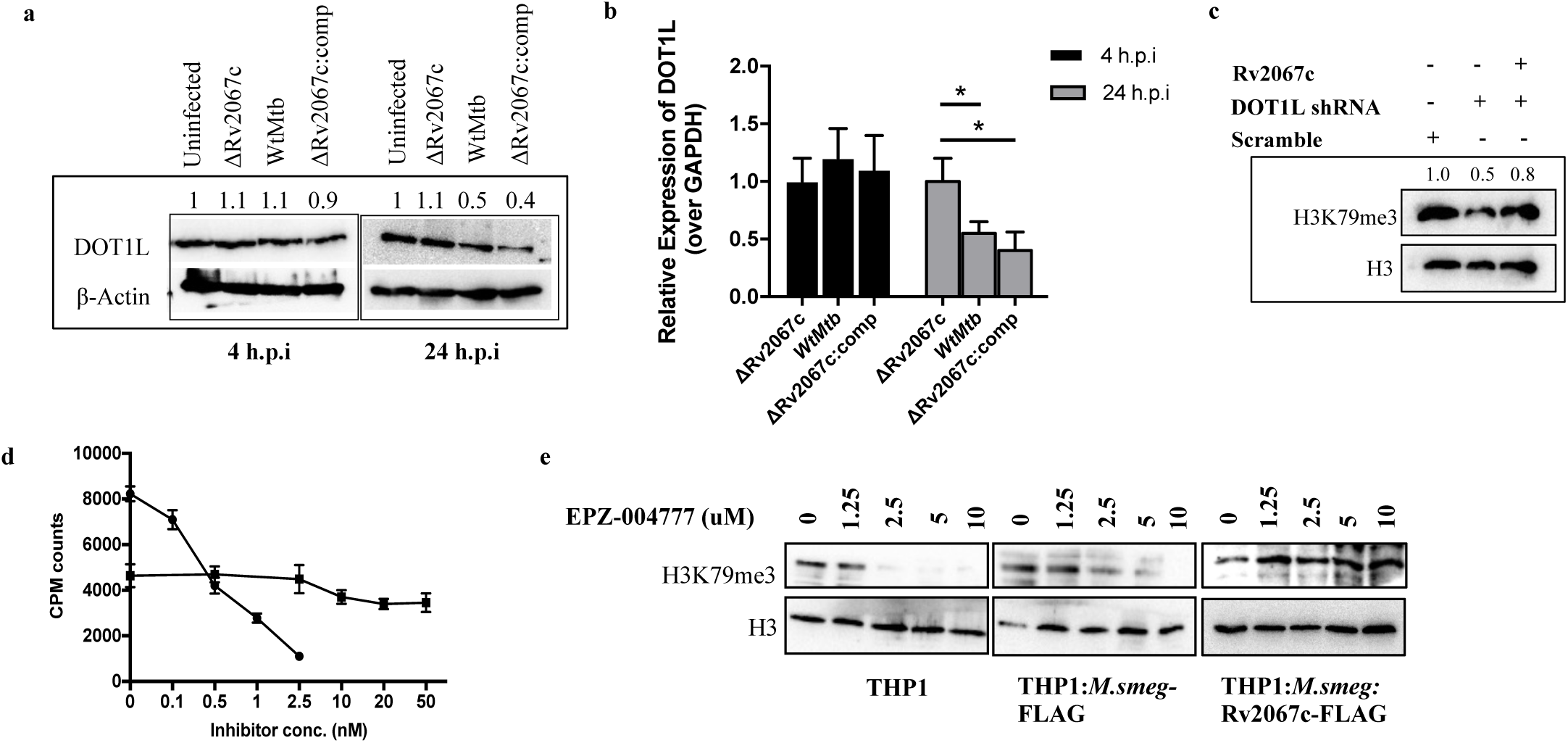
Rv2067c methylates free H3 and modulates DOT1L expression. **a,** Immunoblot depicts level of DOT1L in THP1 cell lysates 4 and 24 h.p.i with various *Mtb* strains. β-Actin was used as loading control. Lane1: uninfected THP1 macrophages; Lane 2, 3 and 4: THP1 macrophages infected with ΔRv2067c, Wt*Mtb* and ΔRv2067c:comp strains, respectively. **b,** Relative expression of DOT1L in THP1 macrophages 4 (black bars) and 24 (grey bars) h.p.i with ΔRv2067c, Wt*Mtb* and ΔRv2067c:comp with respect to uninfected THP1 macrophages. Levels were normalised against GAPDH. The data plotted is mean of two independent infections. Two technical replicates were used for qRT-PCR. Error bars correspond to s.d and *P-value <0.05 (Student’s t-test).**a,b** Comp stands for complemented **c,** Western blot depicts H3K79me3 levels in cell lysates of HEK293T with following treatment: Lane1: scramble shRNA (control); Lane 2: DOT1L inhibition by shRNA and Lane3: DOT1L inhibition by shRNA along with expression of Rv2067c. H3 was used as loading control. **d,** Graph depicts inhibition of recombinant Rv2067c and DOT1L with small molecule inhibitor EPZ004777. Filled circle and filled square represent scintillation counts for DOT1L and Rv2067c with EPZ004777 inhibitor at indicated concentrations. Each data point represents the mean of two replicates at each specified concentration of compound and the error bars represent s.d. **e,** Immunoblot analysis of H3K79me3 in uninfected THP1 macrophages (left panel), inhibitor treated THP1 macrophages infected with *M.smeg*:FLAG (middle panel) and *M.smeg*:Rv2067c-FLAG (last panel) at indicated concentrations. H3 was kept as loading control. Values above the blot represent quantitation (arbitary units).

To ascertain that the enhanced H3K79me3 mark is due to Rv2067c MTase activity, we inhibited DOT1L in two ways - RNA interference and a small molecule inhibitor. HEK293T transfected with DOT1L shRNA showed reduced DOT1L expression (Extended Data Fig. 7 h and i); a corresponding reduction of H3K79me3 mark was confirmed by immunoblotting (Fig. 7c, centre lane). Cells co-transfected with DOT1L shRNA and pcDNA:Rv2067c showed an increase in H3K79me3 mark, confirming the gain of mark is due to the MTase activity of Rv2067c (Fig. 7c). In the second approach, DOT1L was inhibited using EPZ004777 ^33^ which did not affect Rv2067c activity even at higher concentration (50 nM) (Fig. 7d). Since EPZ004777 specifically inhibits DOT1L and not Rv2067c, macrophages were treated with EPZ004777 prior to infection. While the H3K79me3 levels reduced in uninfected inhibitor treated cells, the methylation levels were restored in Rv2067c infected macrophages, despite of reduction of DOT1L RNA, 24 h.p.i (Fig. 7e and Extended Data Fig. 7j). Together, these two approaches confirm that the increased H3K79me3 is specifically due to Rv2067c.

Next, to understand the basis of DOT1L repression upon *Mtb* infection, we analysed two loci-Peak 1(chr19:2164620-2168335) and Peak 2(chr19:2169309-2181752) for epigenetic marks on DOT1L gene ^34, 35^. Enhanced H3K9me3, H3K27me3 and H3K79me3 and depleted H3K4me3 marks were observed at both the loci in macrophages infected with Wt*Mtb* and ΔRv2067c:comp (Extended Data Fig. 7c-g). The reduced expression of DOT1L upon *Mtb* infection thus could be attributed to the altered methylation marks on the chromatin.

### H3K79me3 mark by Rv2067c alters host gene expression

To identify the genes impacted by H3K79 methylation mark added by Rv2067c, we performed ChIP with H3K79me3 antibody on HEK-DOT1L-KD cells expressing Rv2067c. Among several identified, five loci where H3K79 mark was absent in CD14 monocytes^35^ were chosen for analysis (Extended Data Table 2). Significant enrichment of H3K79me3 mark was observed on three loci- *TMTC1, SESTRIN3* and *NLRC3* (Extended Data Fig. 8b). These genes also showed increased expression in HEK-DOT1L-KD cells expressing Rv2067c indicating H3K79me3 added by Rv2067c is an activating mark (Extended Data Fig. 8c). The enriched mark and enhanced expression was also observed in HEK293T expressing Rv2067c (Extended Data Fig. 8b and c). Further, macrophages infected with ΔRv2067c showed decreased H3K79me3 mark and reduced expression of TMTC1, SESTRIN3 and NLRC3 (Fig. 8a and b) along with a concomitant change in SERCA2 expression (Extended Data Fig. 8d). Restoration of H3K79me3 mark and corresponding enhanced expression when infected with ΔRv2067c:comp (Fig. 8a and b), establish that Rv2067c adds H3K79me3 mark upon *Mtb* infection on TMTC1, SENSTRIN3 and NLRC3 loci. Infection with ΔRv2067c:RxR showed H3K79me3 mark and expression similar to ΔRv2067c (Extended Data Fig. 8e and f).

**Fig. 8:**
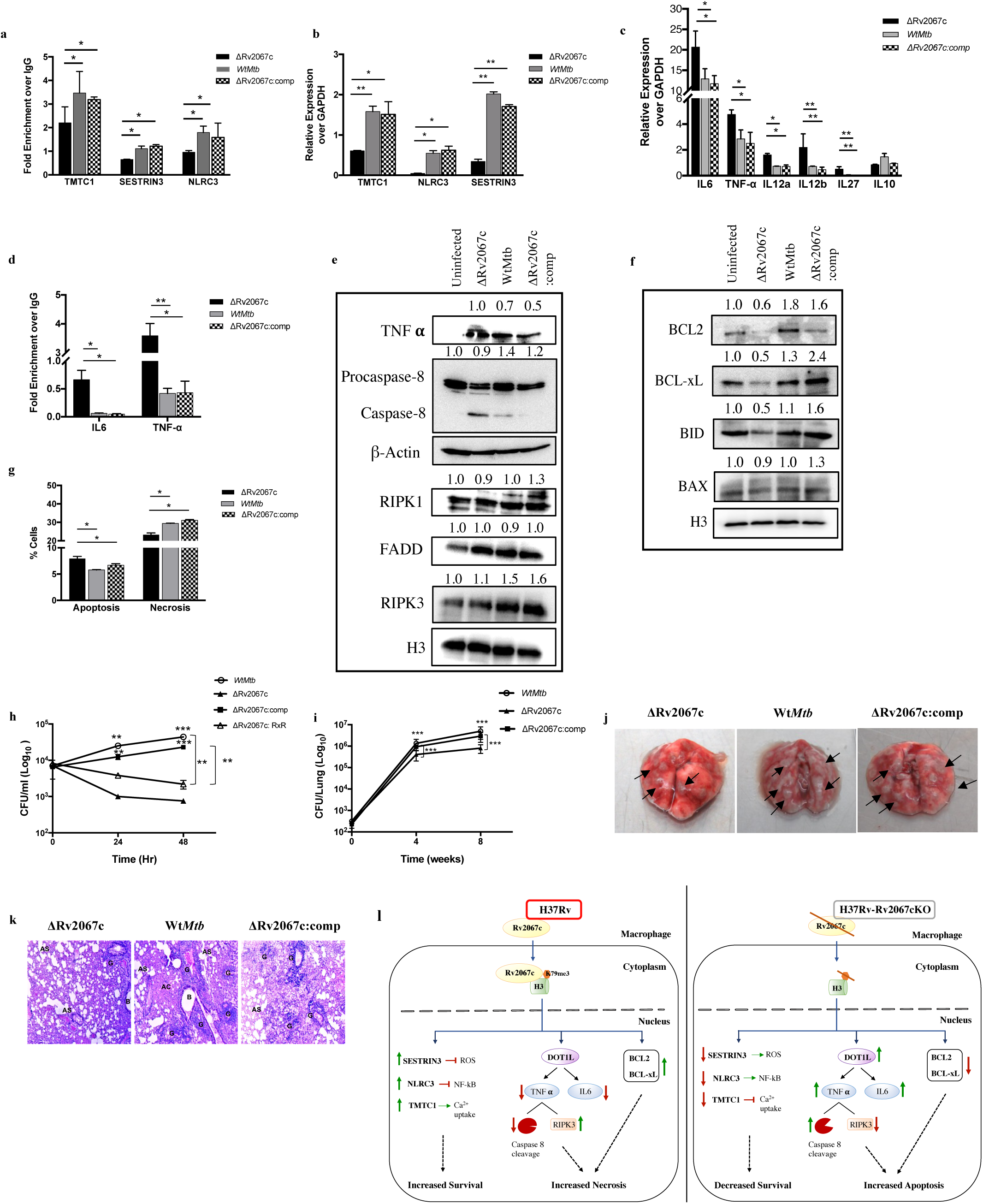
H3K79me3 mark performed by Rv2067c results in gene activation. **a,** Fold enrichment of H3K79me3 mark performed by Rv2067c on specific loci (as indicated) in THP1 macrophages infected with ΔRv2067c (black bars), Wt*Mtb* (grey bars) and ΔRv2067c:comp (checker pattern). Data is shown as fold enrichment over the IgG control. **b,** Relative expression of TMTC1, NLRC3 and SESTRIN3 in THP1 macrophages infected with ΔRv2067c (black bars), Wt*Mtb* (grey bars) and ΔRv2067c:comp (checker pattern) with respect to uninfected THP1 macrophages. Ct values were normalized against GAPDH. **c,** Graph shows relative expression of indicated cytokines in THP1 macrophages infected with ΔRv2067c (black bars), Wt*Mtb* (grey bars) and ΔRv2067c:comp (checker pattern). Ct values were normalized against GAPDH. **d,** Fold enrichment of H3K79me3 mark at the promoter region of IL-6 and TNF-α in THP1 macrophages infected with ΔRv2067c (black bars), Wt*Mtb* (grey bars) and ΔRv2067c:comp (checker pattern). Data is shown as fold enrichment over the IgG control. **a-d**, All ChIP and qRT-PCR data is representative of two independent infections. For qRT-PCR, two technical replicates were kept for each sample. The data plotted is mean and the error bars represent s.d. **e,** Immunoblot depicts expression of indicated genes in THP1 cell lysates 24 h.p.i with various Mtb strains. Lane1: uninfected THP1 macrophages; Lane 2, 3 and 4: THP1 macrophages infected with ΔRv2067c, WtMtb and ΔRv2067c:comp strains respectively. H3 and β-Actin were used as loading control. **f,** Immunoblots depict expression of apoptotic markers in THP1 cell lysates post 24 hr of infection with various Mtb strains. Lane1: uninfected THP1 macrophages; Lane 2, 3 and 4: THP1 macrophages infected with ΔRv2067c, WtMtb and ΔRv2067c:comp strains respectively. H3 was used as loading control.**e and f,** Values above the blot represent quantitation (arbitary units). **g,** Graph depicts percentage of apoptotic and necrotic macrophages analyzed by flow cytometry upon infection with ΔRv2067c (black bars), WtMtb (grey bars) and ΔRv2067c:comp (checker pattern). **h,** Survival of intracellular bacilli post 24 and 48hr of infection in THP1 macrophages with ΔRv2067c (filled triangle), WtMtb (open circle), ΔRv2067c:RXR (open triangle) and ΔRv2067c:comp (filled square). The data is CFU average of three independent infection per time point. Three technical replicates were kept during each infection. Error bars correspond to s.d. CFU, colony forming units. **i,** Bacillary burden in lungs of mice infected with ΔRv2067c (filled triangle), WtMtb (open circle) and ΔRv2067c:comp (filled square), 4- and 8-week post infection (n = 6 animals per group per time point, two technical replicates for plating were kept). CFU, colony forming units. **j,** Gross pathology of the lungs of BALB/c mice infected with ΔRv2067c, WtMtb and ΔRv2067c:comp harvested after 8 weeks of infection. **k,** Haematoxylin and eosin–stained lung sections post 8 week of infection in mice with ΔRv2067c, WtMtb and ΔRv2067c:comp.The pathology sections show granuloma (G), alveolar space (AS), Alveolar consolidation (AC) and bronchial lumen (B). *, P-value <0.05; **,P-value <0.01; ***,P-value <0.001 (Student’s t-test).

### DOT1L repression by Rv2067c impacts cytokine response and cell death pathways

Reduced DOT1L expression in macrophages infected with *Mtb* expressing Rv2067c indicated another role for Rv2067c in manipulating host epigenetic circuitry. Recruitment of DOT1L to the promoter of specific cytokines results in addition of H3K79me2/3 mark and facilitating their expression^36^. Hence, reduced DOT1L due to Rv2067 action could lead to the downregulation of DOT1L responsive cytokines. Transcriptomic analysis of THP1 macrophages infected with *M.smeg*:Rv2067c-FLAG and *M.smeg* (Extended Data Fig. 8g and h) showed 3497 and 1444 genes were significantly upregulated and downregulated respectively (Fold Change ≥ 1.5). (Source data Table1). Amongst these innate immunity, cytokine activity, chemokine related signalling pathway genes were downregulated^37^ (enrichment score >5 and p-value <0.05) (Extended Data Fig. 8i). Expression of pro-inflammatory cytokines IL6, TNF-α, subunits of IL12 – IL12A and IL12B along with DOT1L were downregulated (Extended Data Fig. 8j). Importantly, reduced expression of these cytokines is seen upon Mtb infection (Wt*Mtb* and ΔRv2067c:comp), 24 h.p.i (Fig. 8c). Further, promoters of IL-6 and TNF-α showed reduction in H3K79me3 mark in macrophages infected with Wt*Mtb* and ΔRv2067c:comp and not with ΔRv2067c (Fig. 8d).

TNF-α is known to induce programmed necrosis as well as caspase-8 mediated apoptosis upon *Mtb* infection^38, 39^.We observed increased cleavage of caspase-8 along with decreased RIPK3 in macrophages infected with ΔRv2067c in comparison to Wt*Mtb* and ΔRv2067c: comp (Fig. 8e). RIPK1 and FADD expression did not alter (Fig. 8e). Notably, anti-apoptotic markers BCL2 and BCL-xL were downregulated in ΔRv2067c but not in Wt*Mtb* and ΔRv2067c:comp infected macrophages. BID (uncleaved) decreased and BAX levels did not alter in macrophages infected with ΔRv2067c (Fig. 8f). Flow cytometry analysis showed higher percentage of macrophages infected with Wt*Mtb* and ΔRv2067c:comp undergo necrosis (Fig. 8g and Extended Data Fig. 8k), suggesting that Rv2067c restricts host mediated apoptosis and may promote necrosis.

Next, we investigated the role of Rv2067c in enhancing the survival of intracellular bacilli upon infection. Both ΔRv2067c and ΔRv2067c:RxR had significantly reduced bacterial survival 24 and 48 h.p.i in macrophages compared to Wt*Mtb* and ΔRv2067c:comp (Fig. 8h). Survival advantage was also observed for *M.smeg*:Rv2067c-FLAG in macrophages (Extended Data Fig. 8l). Notably, bacterial burden of ΔRv2067c reduced in lungs of BALB/c mice 8 weeks p.i when compared to Wt*Mtb and* ΔRv2067c:comp (Fig. 8i). The gross and histopathological changes in the lungs of the ΔRv2067c infected mice 8 weeks p.i were comparable with the bacillary load observed (Fig. 8j and k). The extent of pulmonary tissue destruction was lowest in the lungs of mice infected with ΔRv2067c (score 4) relative to Wt*Mtb* (score 11) and ΔRv2067c:comp (score 11) (Extended Data Fig. 8m). Altogether, these results confirm that Rv2067c is a determinant for *Mtb’s* intracellular survival.

## Discussion

Intracellular pathogens engage the host with diverse strategies for infection, survival and multiplication. One emergent strategy is to circumvent and outwit host response by interfering with the host epigenetic machinery. We have uncovered mechanisms by which a secreted MTase Rv2067c of *Mtb* alters host epigenome ensuring intracellular survival of the pathogen. Rv2067c adds H3K79me3 mark on H3 upon *Mtb* infection, to alter host gene expression, including repression of the host MTase DOT1L. In doing so it pre-empts DOT1L catalysed H3K79me3 in the nucleosomal context. Consequently, DOT1L mediated proinflammatory cytokines and apoptotic pathways are attenuated. By adding H3K79me3 mark on loci where DOT1L specific mark is absent, Rv2067c activates the expression of genes involved in countering host response to pathogen. A combination of these actions by MTase steers the host signalling events towards necrosis.

Intracellular pathogens have adopted successful strategies to perturb host epigenetic pathways providing glimpses into underlying molecular mechanisms^40–43^. The MTase RomA of *Legionella pneumophila* methylates H3K14 in the octamer or nucleosome to impact the expression of host innate immunity genes^40^. *Mtb* MTase Rv1988 adds H3R42me2 mark to repress NOS2, NOXA1, NOX1 and NOX2, but the context of methylation (free or nucleosomal) and downstream events remain to be investigated^42^. Eis, a GNAT family acetyltransferase of *Mtb*^10^ facilitates evasion of autophagy by increasing IL-10 expression through acetylation of H3 at its promoter to activate Akt/mTOR/p70S6K pathway ^44^. Other *Mtb* virulence factors also carry out host epigenetic reprogramming. Secretory protein ESAT-6 remodels the host chromatin and inhibits interferon (IFN)-γ-induced type I as well as type IV CIITA^45^. However, none of these seem to employ multiple approaches to alter host cellular signalling described for Rv2067c here. By adding H3K79me3 mark on TMTC1, SESTRIN3 and NLRC3, which were devoid of H3K79me3 mark before *Mtb* infection, Rv2067c enhances their expression. Upregulation of SESTRIN3 results in scavenging ROS after primary ROS accumulation, thereby delaying apoptosis following *Mtb* infection^46^. NLRC3 negatively regulates NF-kB and CD4^+^ T cell response suppressing innate immunity, thereby promoting *Mtb* survival^47^. Activation of NF-kB in macrophages infected with Rv2067c transposon mutant indicated the role of Rv2067c in suppressing NF-kB^48^, supporting our findings. TMTC1, an endoplasmic reticulum integral membrane protein influences calcium homeostasis by interacting with SERCA2 and steers the cell towards necrotic pathway^38^. Thus, it is apparent that H3K79me3 mark added by Rv2067c enhances expression of genes involved in overcoming host response to infection. In parallel, by repressing DOT1L and the consequent reduction in DOT1L specific H3K79me3 mark, Rv2067c plays a role in reduced expression of the IL-6 and TNF-α. Recruitment of DOT1L to the proximal promoter of IL6 and addition of H3K79me2/3 mark facilitates its transcription activation^36^. We show that by regulating TNF-α expression, Rv2067c reduces caspase 8 cleavage along with upregulation of RIPK3 in macrophages infected with *Mtb.*Concomitantly, expression of anti-apoptotic markers BCL-2 and BCL-xL is enhanced. Following *Mtb* infection, Bcl-xL and RIPK3 are critical for preventing caspase 8 mediated apoptosis and induction of necrosis in the mitochondria of macrophages ^49, 50^. Thus, inhibition of caspase 8 mediated apoptosis by the MTase and parallel promotion of expression of anti-apoptotic genes and RIPK3 benefits the pathogen’s intracellular survival. Infection with pathogens can trigger varied levels of TNF-α expression^51^. Although intracellular TNF-α concentration can determine outcome of *Mtb* infection ^38, 52^, Rv2067c action involving several downstream signalling events enable the pathogen to overcome host response. Hence, multiple strategies orchestrated by *Mtb* through Rv2067c, appear to be a fail-safe mechanism to ensure pathogen’s success (Fig. 8l).

Although the host MTase DOT1L and Rv2067c trimethylate H3K79, their structures are strikingly different. Large monomeric DOT1L has structural elements tailored and optimized to recognize H3 in the NCP, while the architecture of dimeric Rv2067c enables it to recognise free H3 and preclude NCP binding. Being a regulator of host innate immune defence, DOT1L function on nucleosomal H3 upon pathogen infection would enable immediate activation of pro-inflammatory pathways for bacterial clearance. Thus, despite methylating same target as DOT1L, *Mtb’s* epigenetic writer by its distinct structure warrants temporal and spatial separation in its action to reprogram cellular response. Although a few host enzymes post-translationally modify newly synthesised H3 and H4^53, 54^, the methylation of cytosolic H3 by Rv2067c appears to be clever strategy of *Mtb* while encountering the host. By intercepting nucleosomal methylation of H3K79me3 and pre-emptive cytosolic methylation of H3, it provides pathogen multiple options to manipulate downstream pathways. Localization of Rv2067c in the nucleus suggests the possibility of yet to be identified substrates of the MTase and its additional role in epigenetic manoeuvring.

Finally, DOT1L is a versatile MTase participating in various cellular functions in diverse tissues regulating effective humoral immune response, CD4^+^ and CD8^+^ T cell differentiation, embryonic development, cell cycle progression, somatic reprogramming and DNA damage^55–57^. Notably, DOT1L also affects virus multiplication by enhancing the antiviral response^58^ and limits nematode multiplication by controlling CD4^+^ T cell activation^59^. Inhibition or silencing of DOT1L during viral infection resulted in decreased nuclear translocation of NF-kB ^58^. Thus as a regulator of innate immune response, DOT1L seems to be an integral component of host defence arsenal against a variety of infections. While the present study shows the effect of Rv2067c in macrophages, it remains to be seen how the MTase impacts DOT1L function and host signalling in other cells which are also targets for *Mtb* infection. Given its multiple roles in promoting *Mtb* survival, Rv2067c is yet another virulence determinant of the pathogen.

## Materials and Methods

### Growth Conditions

*E. coli* cultures were grown in Luria-Bertani (LB) medium (BD Biosciences). *M.smegmatis (M.smeg)* was cultured in 7H9 media (BD Biosciences) and *Mtb* strains were cultured in 7H9 media supplemented with 1X OADC, 0.1% tween and 0.2% glycerol. When required, the culture medium was supplemented with hygromycin (50 µg/ml for mycobacteria, 150 µg/ ml for *E.coli*), kanamycin (25 µg /ml for mycobacteria and 50 µg /ml for *E.coli*), and ampicillin (100 µg/ ml).

### Crystallization of Rv2067c

Crystallization trails were conducted by sitting drop vapor diffusion method at 22°C, with each drop containing 0.8 µL of tagless protein (20 mg/ml) and 0.8 µL of crystallization solution. Rod-shaped crystals were obtained in less than 24 hr, in conditions (from crystallization screen Morpheus, Molecular Dimensions, UK) containing buffer 1 (0.1 M imidazole/MES acid, pH 6.5) or buffer 2 (0.1 M sodium HEPES/MOPS acid, pH 7.5) with precipitant mix (20% w/v polyethylene glycol 500* monomethyl ether + 10% w/v polyethylene glycol 20,000) and additive mix NPS (0.03 M457 sodium nitrate + 0.03 M sodium phosphate dibasic + 0.03 M ammonium sulfate) or halogens (0.03 M sodium fluoride + 0.03 M sodium bromide + 0.03 M sodium iodide) or carboxylic acids (0.02 M sodium formate + 0.02 M ammonium acetate + 0.02 M sodium citrate tribasic dihydrate + 0.02 M potassium sodium tartrate tetrahydrate + 0.02 M Sodium oxamate). For experimental phasing, crystals were either grown in the presence of or soaked in mother liquor containing sodium iodide (NaI) or 5-amino-2,4,6-triiodoisophthalic acid (I3C).

### Data collection, processing, and structure determination

Diffraction data were collected at MX beamline ID29, European Synchrotron Radiation Facility (ESRF), Grenoble. For each dataset, 3600 images were collected with 0.1° oscillation at X-ray wavelength of 1.7000 Å (anomalous) or 1.0723 Å (native). All the datasets were collected at cryogenic temperature (100 K). Diffraction images were processed using the XDSAPP3.0^60^. The best anomalous signal was obtained from crystals grown in buffer 1 with precipitant mix and NPS, and soaked in a mother liquor containing 150 mM NaI for 5 min. Anomalous dataset was processed to 3.25 Å resolution. A better resolution (2.40 Å) dataset with weak anomalous signal from crystals grown in the presence of 10 mM I3C was processed as a native dataset. No native dataset diffracted better than 2.40 Å. Both the datasets belonged to the P4_1_2_1_2 space group. Initial phases were obtained by single anomalous dispersion (SAD) method with iodine as a marker atom, using CRANK2^61^ from CCP4i2 suite^62^. A near complete model was built using Buccaneer^63^. The 2.40 Å resolution structure was determined by PHASER’s molecular replacement (MR) module^64^ using phased structure as MR search model. Alternate cycles of model building and refinement were carried out using Coot^65^ and Refmac5 ^66^, respectively. Model quality was assessed using MolProbity^67^.

### Cell culture and macrophage infection

THP1 monocytes were cultured in RPMI medium supplemented with 10% fetal bovine serum at 37°C and 5% CO_2_. Monocytes were differentiated for 18 hr with 10 ng of PMA and after 24 hr of recovery in PMA-free RPMI medium, infection was performed. Differentiated THP1 macrophages are referred as macrophages in the manuscript.

For infection, bacterial strains were grown up to an OD_600_ of 0.6-0.8, washed once with PBS and re-suspended in medium. Infections were carried out at 1:10 MOI for *M.smeg* and 1:5 MOI for *Mtb* strains respectively, unless otherwise mentioned. After 4 hr of infection, gentamycin (50ug/ul) was added to kill any extracellular bacteria. Media was changed after 1 hr of gentamycin treatment and cells were incubated for required time.

Cells were harvested after washing with ice-cold phosphate-buffered saline (PBS) and lysed in 1X RIPA buffer (50 mm Tris-HCl, pH 7.4, 1% Nonidet P-40, 0.25% sodium deoxycholate, 150 mm NaCl, 1 mm EDTA). Cell lysates were spun at 12,000 rpm for 20 mins at 4°C. Supernatant was collected and 50-100 ug was loaded on SDS-PAGE for western analysis. RNA was isolated from cells using TRI reagent (Sigma). For qRT-PCR, 1-2 ug of RNA was converted to cDNA using reverse transcriptase (Applied Biosystems).

### Generation of Rv2067c strains in *Mtb*

To generate a knockout of Rv2067c by homologous recombination, 3’(−953 to 53 bp) and the 5’ regions (+1180 to + 2209 bp) of Rv2067c were amplified from the *Mtb* H37Rv genomic DNA with specific primers. 5’ flanking region was cloned into SpeI and SwaI digested pML523 followed by cloning of 3’ region into PacI and NsiI digested pML523 vector. The complete 4628 bp DNA fragment was PCR amplified and sub-cloned in pRSF-Duet digested with EcoRV digestion. This construct was transformed into wildtype *Mtb* H37Rv (Wt*Mtb*). An internal fragment of the Rv2067c ORF was replaced by a LoxP-GFP:hyg-LoxP cassette. Genomic DNA was isolated from colonies. Knockout was confirmed by PCR and western blotting with Rv2067c antibody. Rv2067c antibody was raised in rabbit and antibody specificity was checked by western analysis (Extended Data Fig. 6c). To unmark the deletion mutant, the pCreSacB plasmid was transformed into deletion mutant. Unmarking of the transformants was confirmed by PCR with hygromycin specific primers. The unmarked strain is referred as ΔRv2067c or knockout in the manuscript. Complemented strains were generated by transforming pST-2KRv2067c and pST-2KRv2067cRxR in ΔRv2067 background. These strains are referred to as ΔRv2067c:comp or complement and ΔRv2067c:RxR respectively.

Over expression strain of Rv2067c in wt*Mtb* H37Rv was generated using inducible pST-KT vector^68^. The gene was cloned between NdeI and HindIII restriction sites. Over expression of Rv2067c in transformants was confirmed with a concentration gradient of ATc followed by immunoblotting with Rv2067c antibody. *Mtb* Rho was kept as a loading control (Extended Data Fig. 5f). The overexpression strain is referred to as Rv2067c:OE in the manuscript.

### Methyltransferase assay

*In vitro* methyltransferase (MTase) assays were carried out by incubating 500ng of methyltransferases Rv2067c or DOT1L with 1–2 μg of recombinant mammalian histones or MtHU and 80μCi ^3^H-SAM (PerkinElmer) in a buffer containing 50 mM Tris-HCl (pH 8.0), 5% glycerol, 5 mM MgCl_2_, 1 mM dithiothreitol (DTT) and 50mM NaCl at 37°C for 30min. Buffer composition for MTase assays with histone octamer and NCPs as substrates was adapted from Min J et al^19^. The reactions were stopped by adding 20% trichloroacetic acid followed by incubation on ice for 1 hr. The mixture was spotted on cellulose nitrate filters (Sartorius) using a vacuum manifold and washed with 20% trichloroacetic acid, water and 95% ethanol, dried at RT, soaked in 3ml of scintillation fluid and counts (CPMs) were measured in a scintillation counter (Perkin Elmer). For autoradiography, the reaction mixture was resolved on a 15% SDS–polyacrylamide gel electrophoresis (PAGE), electro-blotted on polyvinylidene fluoride (PVDF) membrane (GE Healthcare) and incubated for 1 hr in 1% sodium salicylate solution (Sigma). The membrane was exposed to an X-ray film at −80 °C. Salt extraction of histones was performed from THP1 cells as described previously^69^. Extracted histones were sequentially dialyzed to reduce NaCl concentration to 200mM. 20 ul of extracted histones were used as substrate for MTase using SAM as methyl donor. MTase assays with unmodified peptides was carried out using 500ng of peptide (AnaSpec) as substrate and Rv2067c as MTase. The reactions were incubated at 37°C for 30min, spotted on nitrocellulose membrane and immunoblotted with H3K79me3 antibody.

### Immunofluorescence

For immunofluorescence, 0.2×10^6^ macrophages were seeded and infected with Rv2067c:OE, ΔRv2067c, *M.smeg*:FLAG and *M.smeg*:Rv2067c-FLAG. Cells were washed with PBS 24 hr post infection (h.p.i), fixed using 4% paraformaldehyde, permeabilized with 0.1% Triton X-100 and blocked with 2% BSA. FLAG or Rv2067c antibodies were used as primary antibodies and alexa Fluor conjugated antibodies were used as secondary antibodies. Nuclei were stained with diamidino-2-phenylindole dye (DAPI) and the cells examined by confocal microscopy.

### Preparation of culture filtrate

Electrocompetent *M.smeg* cells were transformed with pVV16:Rv2067c-FLAG plasmid. *M.smeg* expressing Rv2067c-FLAG is referred as *M.smeg:*Rv2067c-FLAG. 1% primary culture was inoculated in 100ml of modified Sauton’s media and at OD_600_ of 0.6–0.8 cells were harvested by centrifugation at 8,000 r.p.m. for 30 min. The supernatant was passed through 0.45 μm filter to remove cells and concentrated using 30 kDa cut off centricon (Millipore). Cell pellet was resuspended in PBS and cell lysate was prepared. Secretion of Rv2067c from Wt*Mtb* strain was detected by concentrating 300ml of culture filtrate followed by TCA precipitation.

### Inhibition of DOT1L

The protocol for MTase assay with DOT1L inhibitor was adapted from Daigle, S. R. et al^33^. Briefly, for *in vitro* assay, inhibitor EPZ004777 at a concentration upto 50nM was incubated for 15 min with 500ng of Rv2067c in MTase assay buffer. Post incubation, 1ul of tritiated SAM and 2ug of substrate were added to each of the reaction. Reactions were incubated for additional 30 min at 37°C and radioactive counts (CPMs) were measured in a scintillation counter. A positive control of 2ug of reconstituted NCPs as substrate, recombinant DOT1L as MTase along with the inhibitor was kept. For assessment of EPZ004777 on Rv2067c upon infection, 3×10^6^ macrophages were seeded in 6 well plate and exposed to varying concentration of EPZ004777. 12 hr after exposure to the inhibitor, cells were infected with *M.smeg*:FLAG and *M.smeg*:Rv2067c-FLAG. Cell lysates were probed with H3K79me3 antibody 24 h.p.i. Histone H3 served as a loading control.

For RNA interference, shRNA construct targeting DOT1L was transfected into HEK293T cells and puromycin selection was applied for 24 hr. RNA was isolated followed by qRT PCR to confirm DOT1L inhibition. DOT1L silenced HEK cells were transfected with pcDNA:Rv2067c puromycin construct and maintained under puromycin selection for another 24 hr. Cell lysates were prepared and immunoblotted with H3K79me3 antibody.

### Identification of genomic loci with H3K79me3 mark performed by Rv2067c

To identify H3K79me mark specifically added by Rv2067c, HEK293T cells were co-transfected with DOT1L siRNA and pcDNA:Rv2067c-SFB using Lipofectamine 2000 (HEK-DOT1L-KD:Rv2067c). DOT1L siRNA (AM16708) and scrambled siRNA were procured from Thermo Fisher Scientific. HEK293T cells co-transfected with scrambled siRNA and pcDNA-FLAG were kept as control (HEK-scr:pcDNA). Knockdown of DOT1L in HEK cells by siRNA was confirmed by western analysis (Extended Data Fig. 8a).

The transfected cells were subjected to ChIP with H3K79me3 antibody as described by Yaseen et al.^42^ with minor modifications. The enriched DNA was end-repaired using DNA polymerase I, Klenow fragment (NEB) and adaptor ligated with annealed adaptors (Nimblegen Systems). Next, the enriched DNA was amplified by PCR using LK102 adaptor as primer and resolved on 1.5% agarose gel. The amplified product was excised, eluted and cloned in pBSK vector. After transformation, individual colonies were screened using BamHI and HindIII and positive clones were sequenced. The sequence obtained was aligned using Blat on UCSC genome browser and co-ordinates with enriched H3K79me3 mark were identified.

### RNA sequencing

THP1 macrophages were infected with *M.smeg*-FLAG and *M.smeg*:Rv2067c-FLAG. 4 and 24 h.p.i, macrophages were washed with PBS and RNA was isolated using RNeasy kit (Qiagen). For transcriptomic analysis, sequencing libraries were prepared using NEBNext® UltraTM II RNA Library Prep Kit for Illumina® following the manufacturer’s instructions. Briefly, 800 ng of total RNA was used as input for poly(A) mRNA enrichment followed by fragmentation and reverse transcription to generate cDNA. Hairpin adapter was ligated to fragmented double strand cDNA and USER enzyme was used to cleave the hairpin structure. Ampure beads were used to purify adapter ligated fragments and the purified product was amplified using Illumina Multiplex Adapter primers to generate sequencing library with barcodes for each sample. RNA sequence data was generated using Illumina HiSeq. RNAseq was carried out by Clevergene, India.

### Mass spectrometry

MTase assay was carried out with recombinant histone H3 as a substrate and Rv2067c as MTase. The reaction was incubated overnight at 37 °C. Sample was resolved on 12% SDS– PAGE, gel slice was excised and subjected to MS/MS analysis along with recombinant H3 as a control. For detection of Rv2067c in the culture filtrate, concentrated culture filtrate was resolved on 12% SDS–polyacrylamide gel electrophoresis for 15min. 1cm of gel piece was excised and analyzed by mass spectrometry (Taplin Mass spectrometric Facility, Harvard, USA).

### Affinity pull-down

Rv2067c with N-terminal SFB-tag (S-protein, FLAG, streptavidin-binding peptide) was cloned in pcDNA3.1(pcDNA:Rv2067cSFB) and transfected into HEK293T cells using Lipofectamine 2000 (Invitrogen). A control of HEK293T cells transfected with pcDNA SFB was kept. After 24 hr, cells were lysed in buffer containing 20 mM Tris-HCl, pH 8.0, 100 mM NaCl, 1 mM EDTA, 0.5% NP-40 and protease inhibitors. Lysates were incubated with pre-equilibrated streptavidin beads (GE Healthcare) for 4 hr, the beads were washed three times with the same buffer, loaded on a 12% SDS–PAGE after resuspending in 6X SDS sample loading buffer and western blotted. The blot was probed with H3 and FLAG antibodies.

For reverse immunoprecipitation (IP), same protocol for transfection and cell lysis was followed. Cell lysates were incubated with H3 antibody coated on protein A/G beads for 4 hr. IgG was used as antibody control. The blot was probed with H3 and FLAG antibodies. Affinity pull down using FLAG antibody on nuclear and cytosolic fractions was carried out using a modified protocol from Loyola A et al^54^.

### Animal Experiments

BALB/c mice were infected by aerosol with 200 bacilli per mouse with Wt*Mtb,* ΔRv2067c and ΔRv2067c:comp. At indicated times p.i, mice were euthanized, lungs were harvested for bacillary load, tissue histopathology and granuloma scoring was performed as described before ^70^. The lungs were homogenized and bacillary load was quantified by plating serial dilutions onto Middlebrook 7H11-OADC agar plates supplemented with lyophilized BBL MGIT PANTA antibiotic mixture (polymyxin B, amphotericin B, nalidixic acid, trimethoprim, and azlocillin, as supplied by BD; USA). Colonies were counted after 3-4 weeks of incubation at 37°C. Animal studies were carried out in strict accordance with the guidelines prescribed by the Committee for the Purpose of Control and Supervision of Experiments on Animals (CPCSEA), Government of India. All efforts were made to minimize the suffering. Experiments were carried out in a biosafety level 3 containment facility and approved by the Institute’s Animal Ethical Committee (IAEC), Indian Institute of Science (IISc), Bangalore, India (Approval number: CAF/Ethics/850/2021).

### Statistical analysis

Statistical significance for comparison between two groups was determined by the student’s t-test using GraphPad Prism software (7.0, 9.0 versions, GraphPad Software, USA) and excel. *, P-value <0.05; **,P-value <0.01; ***,P-value <0.001. Quantitation of western blots was performed using Image J (arbitrary units normalized with the expression of the H3 or β-actin).

For MTase assay, data plotted is mean ± standard deviation for 3 independent experiments. For ChIP and qRT, data shown is mean ± standard deviation of two independent sets of infection. For qRT-PCR, two technical replicates per sample were kept.

## Supporting information

Supplementary Results and methods

## Data availability

### Transcriptomic Data

Raw sequencing data has been deposited to the Short Read Archive (SRA) under the project number PRJNA907927.

### Accession code

Coordinates for Rv2067c-SAM are deposited in the RCSB Protein Data Bank under the accession code 8HKR.

Data Collection for rotation scan was done using the script submitted at https://github.com/Venkat-Dadi/Rotation_Scan.

### Disclosures

The authors declare no competing interests

## Acknowledgements

We thank K.N.Balaji for insightful discussions, S.R. Gangi Setty, N. R. Sundaresan, S.C. Raghavan for providing reagents and Harsh Bansia for initial crystallographic efforts. We thank European Synchrotron Radiation Facility (ESRF), France, for X-ray data collection and D.A. Case, Rutgers University for Amber20 licence. All experiments with *Mtb* H37Rv were carried out in BSL3 facility, Centre for Infectious Disease Research (CIDR), IISc. FACS facility at CIDR, central facility of phosphor imaging and real time PCR at Biological Science Division, Indian Institute of Science (IISc) are acknowledged. PRS is supported by post-doctoral research associateship of Jawaharlal Nehru Centre for Advanced Scientific Research (JNCASR) and VD by senior research fellowship from University Grants Commission(UGC), Government of India.

---

**Extended Data Fig. 1.**
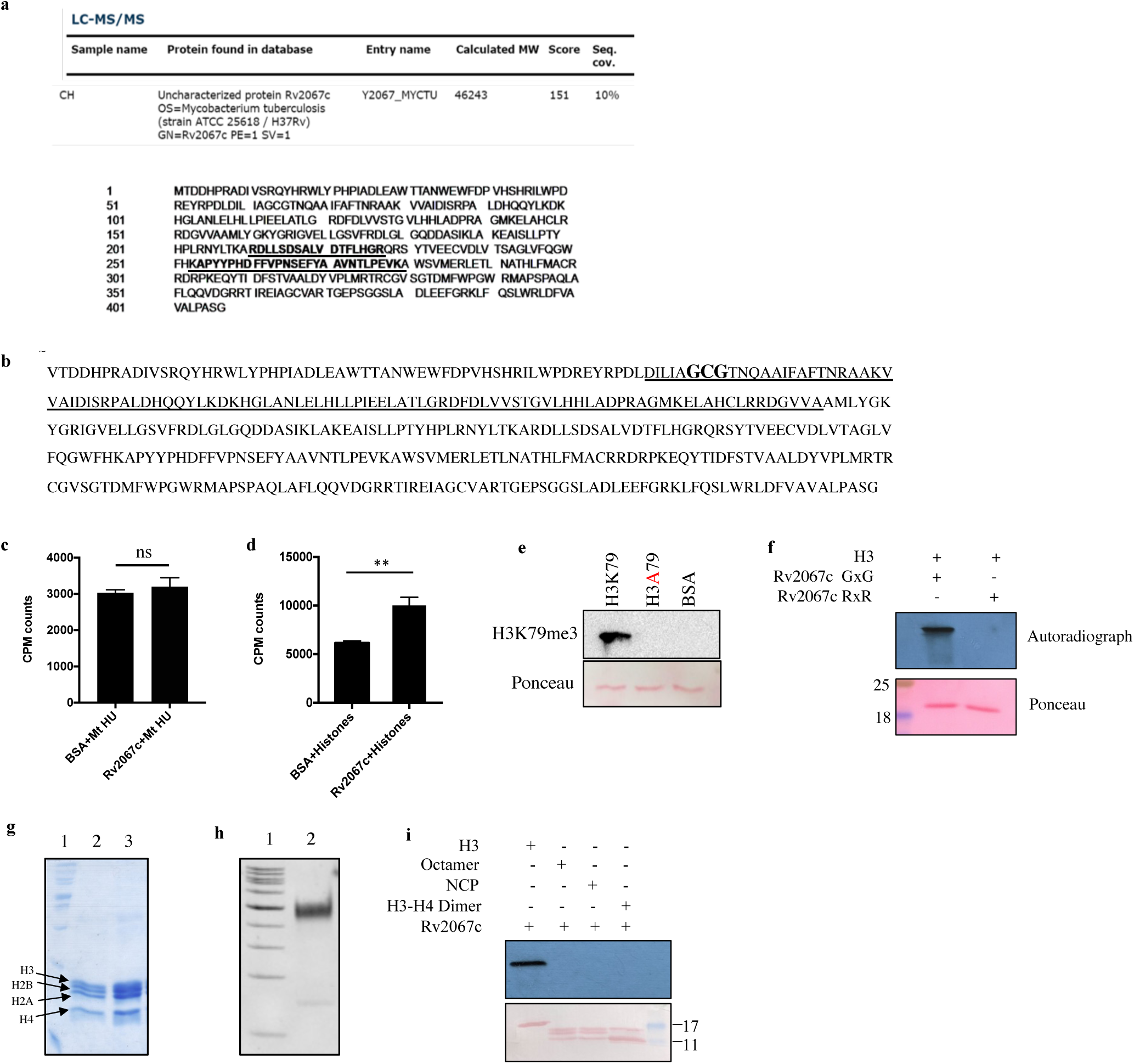
Rv2067c methylates histone H3 at Lysine 79. **a,** Rv2067c was immunoprecipitated from *Mtb* H37Ra cell lysates using affinity purified MtHU antibody. The unique tryptic digests specific to Rv2067c identified by LC-MS/MS are shown in bold and underlined. **b,** Protein sequence of Rv2067c. Putative methyltransferase domain identified using NCBI conserved domain search is underlined, and SAM binding motif is marked in bold. **c and d,** Graph depicts scintillation counts (CPM) for *in vitro* MTase assays with MtHU and salt extracted histones from THP1 monocyte, respectively, as substrates, Rv2067c as MTase and tritiated SAM as a methyl group donor. BSA was kept as negative control. The data plotted for MTase assay is average of three independent experiments. Error bars represent s.d and **,P-value <0.01, ns - not significant (Student’s t-test). **e,** Western blot for *in vitro* MTase assay with recombinant H3 and H3A79 mutant protein as substrate and Rv2067c as MTase. Blot was probed with H3K79me3 antibody. Ponceau staining of blot was used as loading control. **f,** Autoradiograph shows methylation activity of recombinant Rv2067c GxG and Rv2067c RxR with H3 as substrate and tritiated SAM as a methyl group donor. Ponceau staining was used as loading control. **g,** Gel picture depicts reconstituted human histone octamers. Lane 1: protein marker, Lane 2,3: fractions of size exclusion chromatography. **h,** Figure depicts nucleosome preparation from octamers and Widom 601 DNA by micro batch reconstitution method. **i,** Autoradiograph shows methylation activity of Rv2067c with recombinant H3, octamers, NCPs and H3-H4 dimer as substrate and tritiated SAM as a methyl group donor. Ponceau staining was used as loading control.

**Extended Data Fig. 2.**
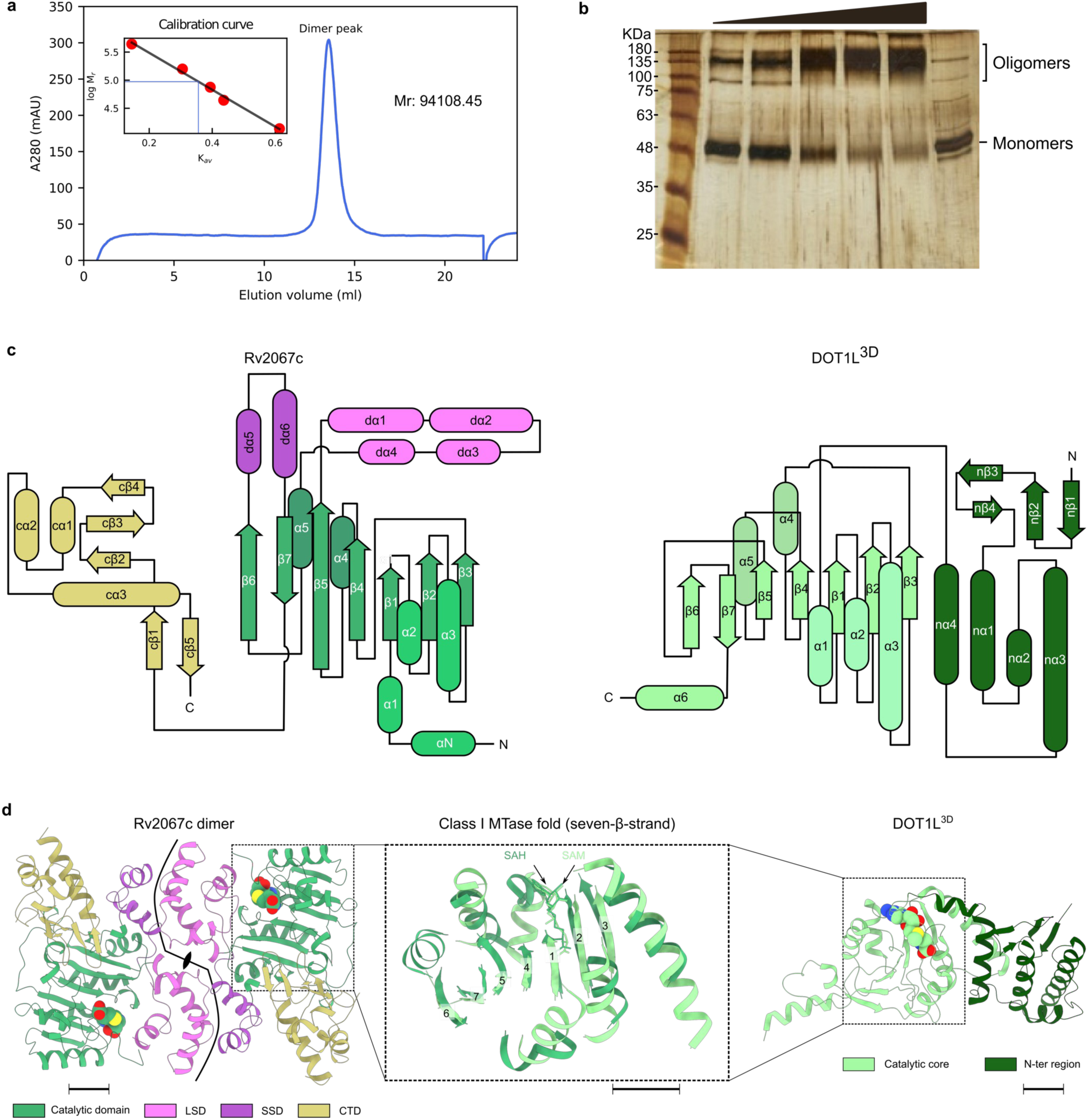
Structural comparison between Rv2067c and DOT1L. **a.** Analytical gel filtration chromatogram showing Rv2067c elution profile. Rv2067c elutes as a dimer with observed Mr 94,108.45 (calculated monomer Mr: 45,930). **b.** Silver-stained PAGE shows the formation of oligomeric Rv2067c upon glutaraldehyde treatment. **c.** Topology diagram of Rv2067c (left panel) and DOT1L^3D^ (right panel), PDB: 1NW3. **d.** Superposition of catalytic cores of Rv2067c and DOT1L. Both share a class I MTase fold (seven β-strand). The β-strands are numbered through 1 to 7.

**Extended Data Fig. 3.**
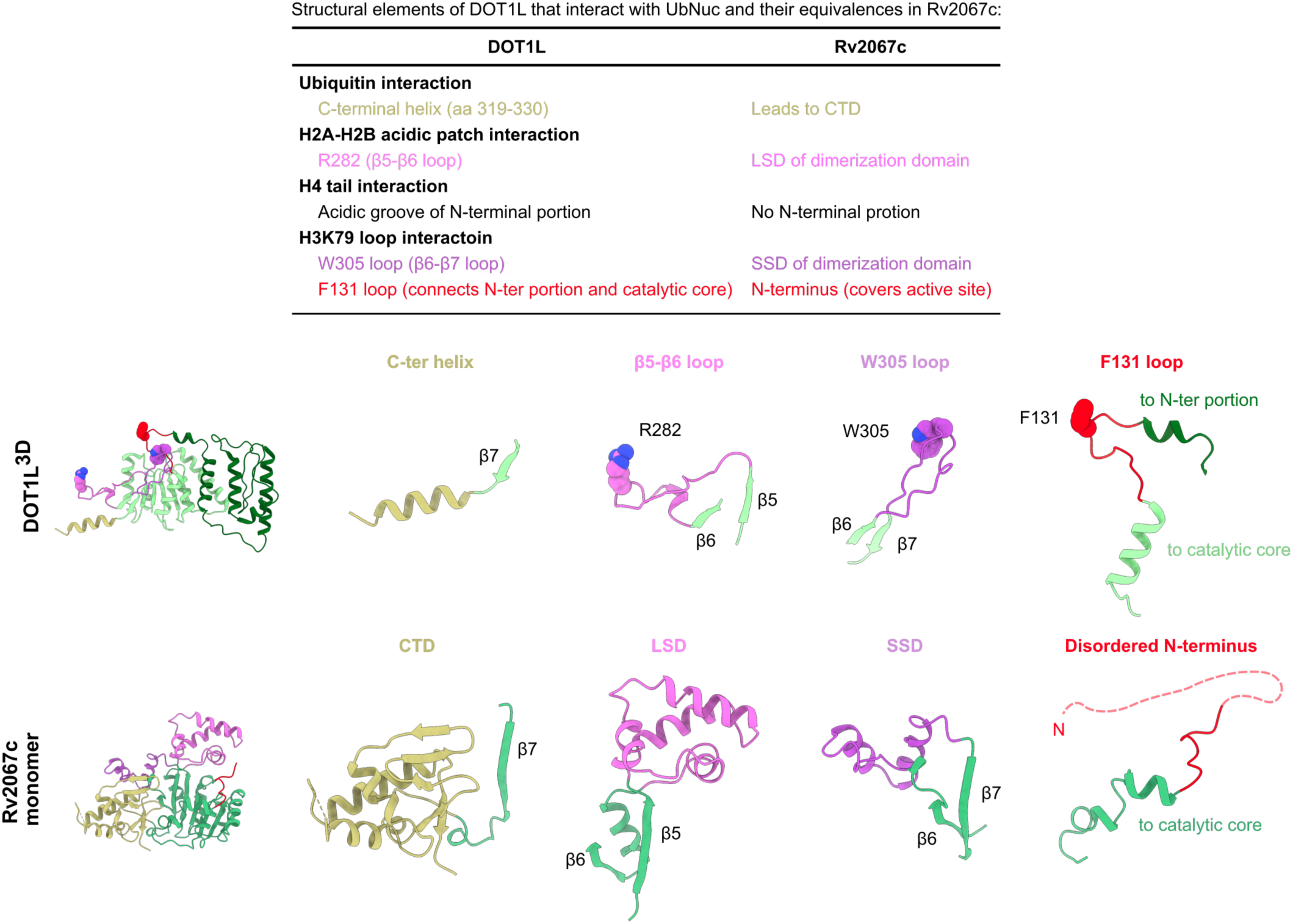
Structural elements of DOT1L essential for nucleosomal H3K79 methylation and their equivalences in Rv2067c.

**Extended Data Fig. 4.**
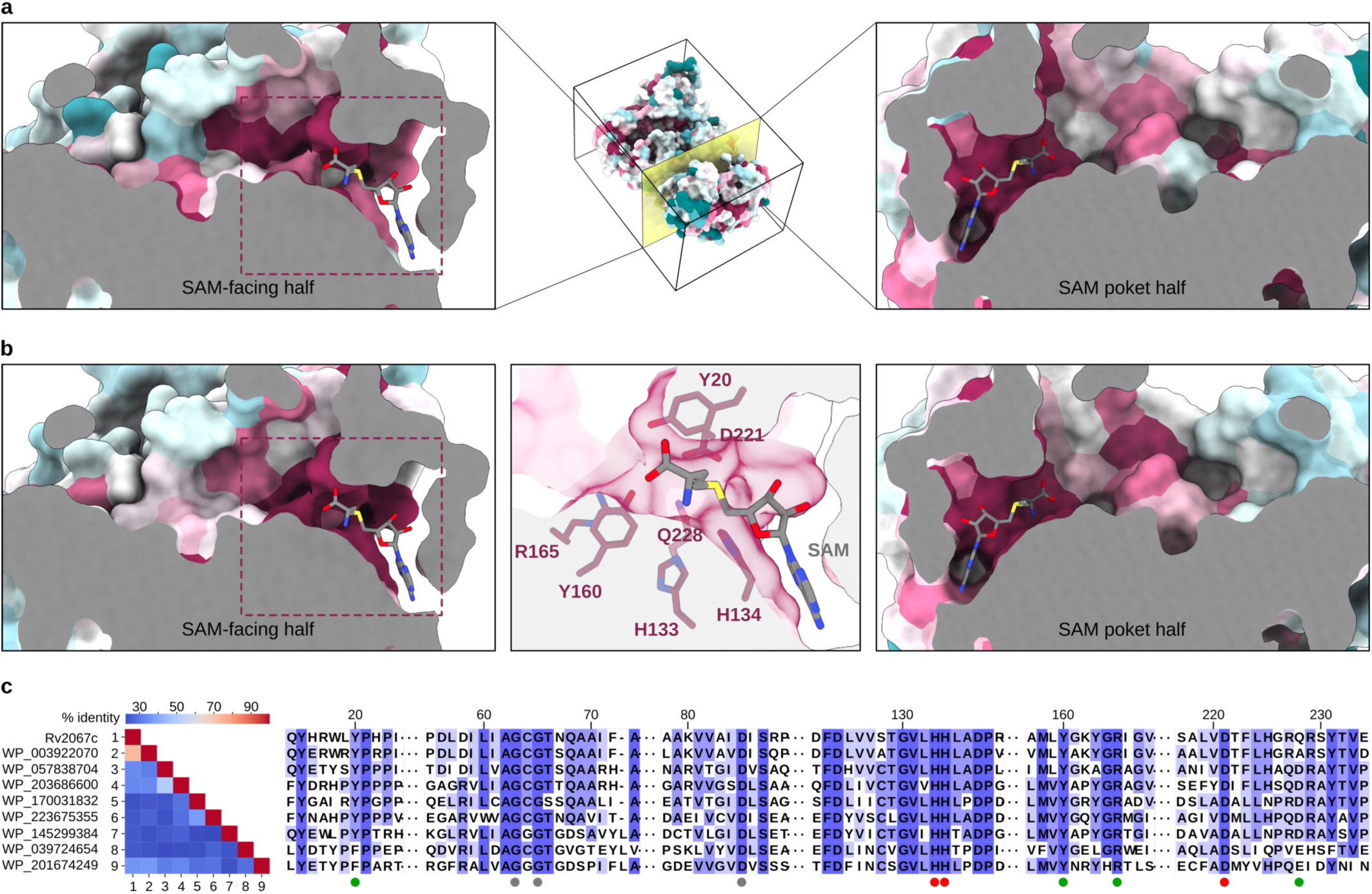
**a** and **b,** Sequence conservation for set A (only mycobacterial species) and set B (all species) was mapped onto the Rv2067c structure using ConSurf. Cross sections along the putative substrate-binding trough (left and right panels) are shown. The region encompassing the SAM is highly conserved compared to the rest of the trough. The putative active site residues (Y20, Y160, R165, D221 and Q228) form a conserved patch opposite to the methyl group of SAM (middle panel of b). **c,** Multiple sequence alignment of a representative sequences from set B. Pairwise sequence identity is shown as matrix. The conserved residues marked with filled circles: green, active site residues (possible substrate lysine binding); gray, SAM-binding (GxG motif and acidic residue); red, putative catalytic residues.

**Extended Data Fig. 5.**
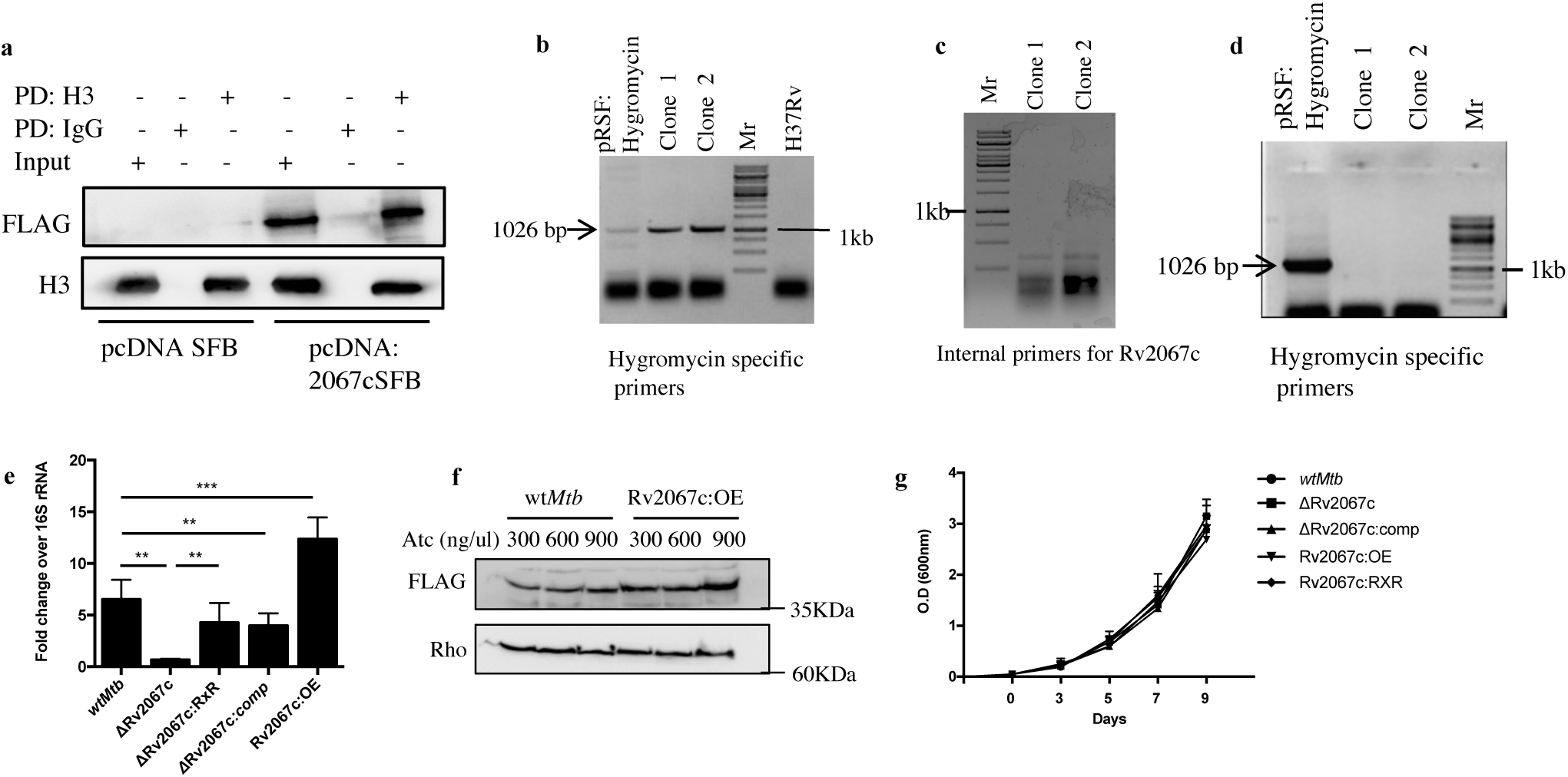
Rv2067c methylates histone H3 at Lysine 79. **a,** Western blot depicts interaction between Rv2067c and H3 by IP performed with H3 antibody on HEK293T transfected with pcDNA: Rv2067cSFB or pcDNA SFB (control) constructs. 5% of whole cell lysate was kept as Input. IP was also performed with IgG as an antibody control and blots were probed with antibodies as indicated. **b and c,** PCR amplification using specific primers (as indicated below the agarose gel) for screening of ΔRv2067c mutant. Gel picture depicts results for two clones. Clones positive for homologous recombination gives a pcr product of 1026 bp with hygromycin specific primers and do not show amplification with Rv2067c specific primers. pRSF:Hygromycin plasmid was kept as positive control for PCR (Lane 1), Mr: 1kb ladder. **d,** Agarose gel depicts unmarking of hygromycin gene in unmarked ΔRv2067c mutant. Unmarked ΔRv2067c positive clones do not give amplification with hygromycin specific primers. pRSF:Hygromycin was kept as positive control for PCR (Lane 1), Mr: 1kb ladder. **e,** Bar graph shows expression of Rv2067c in Wt*Mtb*, ΔRv2067c, ΔRv2067c:RxR, ΔRv2067c:comp, and Rv2067c:OE strains. Levels were normalized against *Mtb* 16S rRNA. The data is representative of two independent RNA isolations from all the strains. Two technical replicates were kept for qRT-PCR. *, P-value <0.05; **,P-value <0.01; ***,P-value <0.001 (Student’s t-test). **f,** Immunoblot depicts over expression of Rv2067c in Rv2067c:OE strain in an Atc dependent manner. Blot was probed with FLAG antibody for Rv2067c. Rho was used as loading control. **g,** Growth curve of different strain generated for the study. The data plotted is mean of two independent experiments and error bars represent s.d.

**Extended Data Fig. 6.**
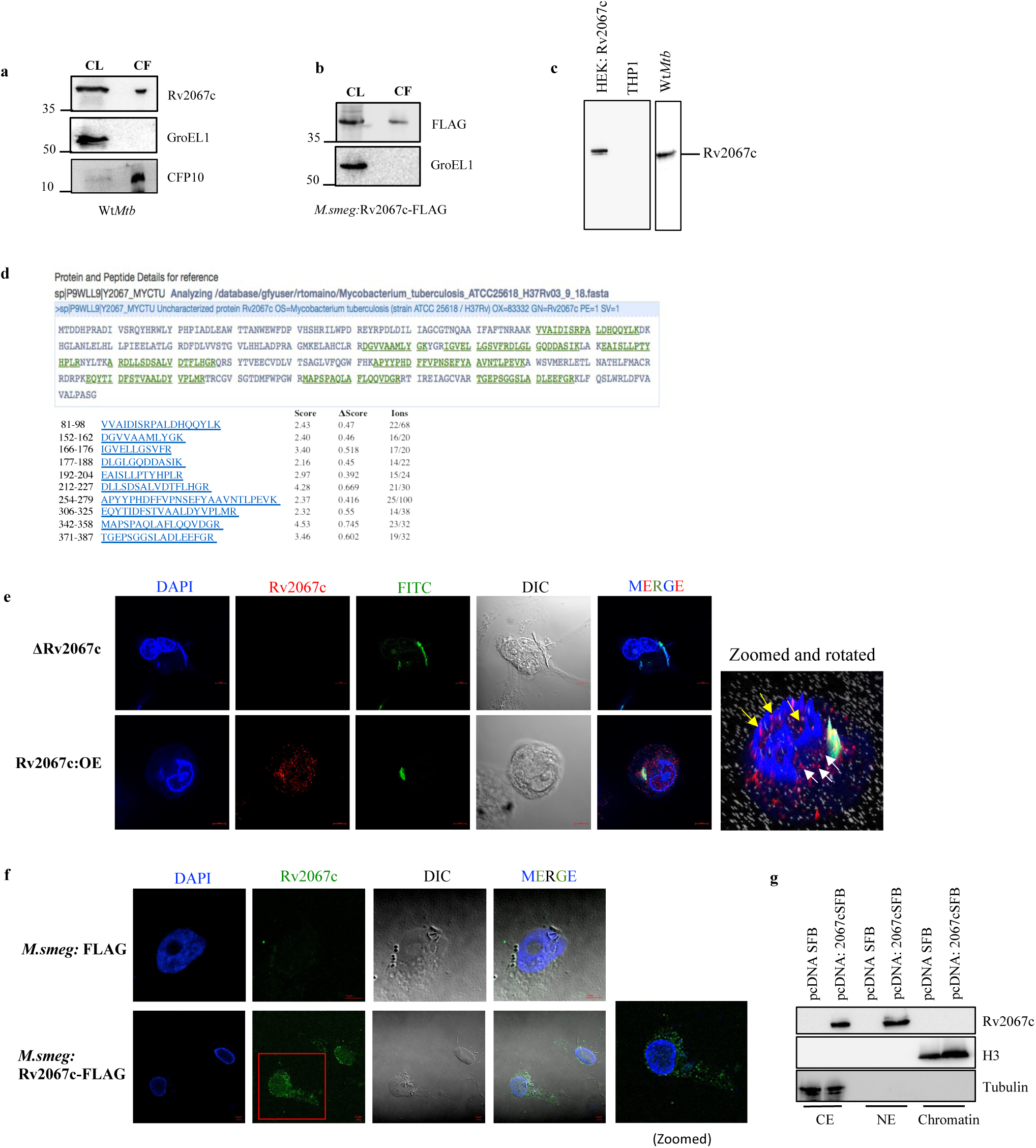
Secretion and localization of Rv2067c. **a,** Western blot for detection of Rv2067c in the culture filtrate (CF) of Wt*Mtb*. Cell lysate (CL) and culture filtrate (CF) were probed with Rv2067c antibody. GroEL1 (a non-secretory protein) antibody was used to examine cell lysis and CFP10 (a secretory protein) was kept as positive control. **b,** Western blot for secretion of Rv2067c from *M.smeg.* Cell (CL) lysate and culture filtrate (CF) of *M.smeg*:Rv2067c-FLAG was probed with FLAG and GroEL1 antibody. **c,** Blot depicts specificity of Rv2067c antibody for cell lysates of HEK293T transfected with pcDNA: Rv2067cSFB, uninfected THP1 and Wt*Mtb*. **d,** Detection of Rv2067c by Mass spectrometry in *M.smeg*:Rv2067c-FLAG culture filtrate. Fragments detected in mass spectrometry are underlined and highlighted in green. **e,** Immunofluorescence analysis of THP1 macrophages infected with FITC labelled ΔRv2067c and Rv2067c:OE (green). Signal for secretory Rv2067c was detected by immunostaining with Rv2067c antibody + Alexa 568 conjugated secondary antibody (red). White and yellow arrows show presence of Rv2067c in the cytoplasm and nucleus respectively. Scale bar=5mm. **f,** Confocal images of THP1 macrophages infected with *M.smeg:*FLAG and *M.smeg*:Rv2067c-FLAG. Signal for Rv2067c was detected by immunostaining with FLAG antibody + Alexa 488 conjugated secondary antibody (green). Inset shows zoomed in view of the region marked by red box. Scale bar=5mm. **g,** Sub-cellular fractions of HEK293T transfected with pcDNA: Rv2067cSFB. Blot was probed with Rv2067c antibody, H3 and tubulin antibody was used as control for chromatin and cytoplasmic extract (CE), respectively. HEK cells transfected with pcDNA:SFB were kept as control.

**Extended Data Fig. 7.**
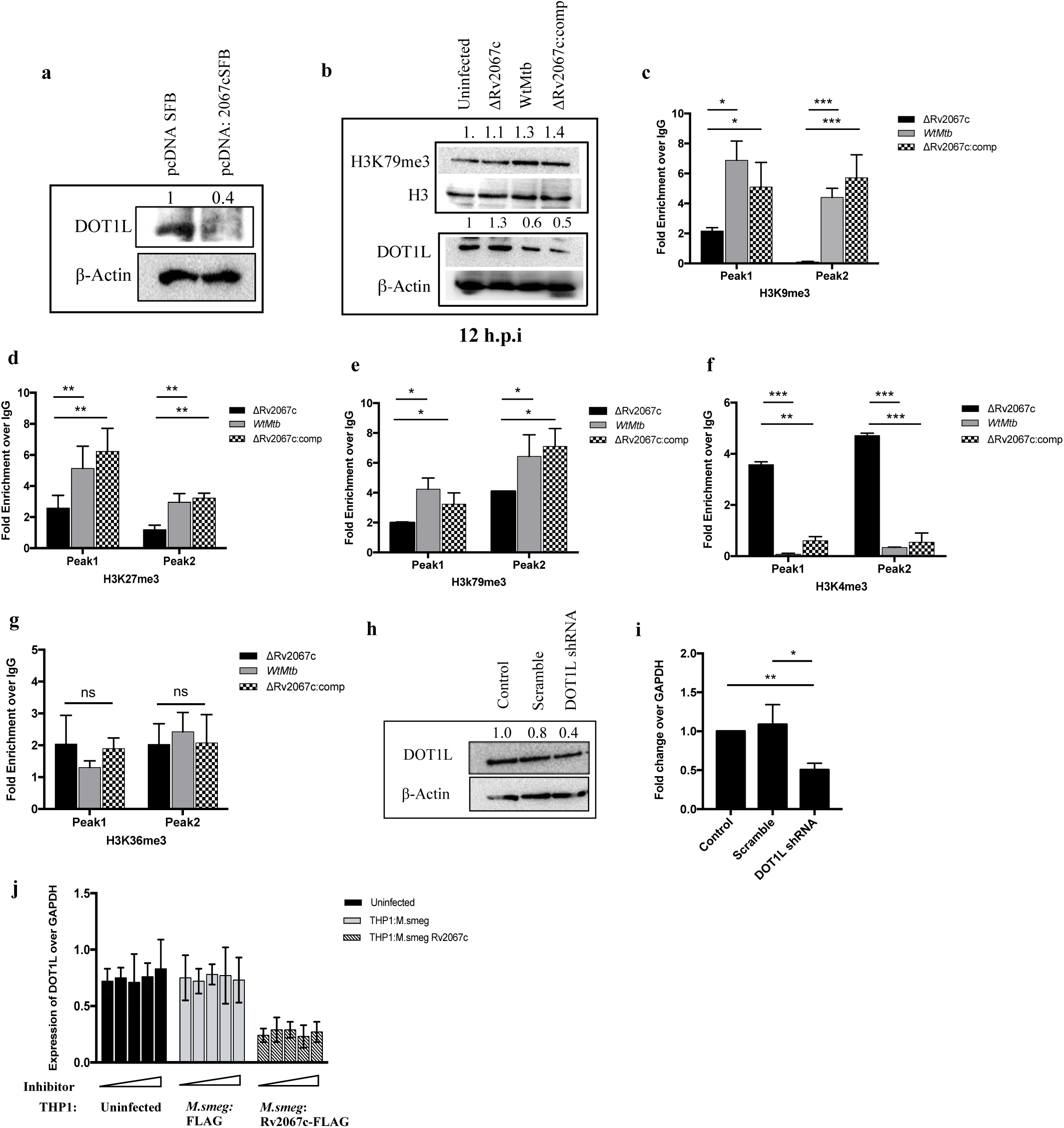
Rv2067c methylates free H3 and modulates DOT1L expression. **a,** DOT1L level in cell lysates of HEK293T transfected with pcDNA: Rv2067cSFB and pcDNA SFB construct. β-Actin was kept as loading control. **b,** Immunoblot depicts level of DOT1L and H3K79me3 in THP1 cell lysates 12 h.p.i with various *Mtb* strains. H3 and β-Actin were used as loading control. Lane1: uninfected THP1 macrophages; Lane 2, 3 and 4: THP1 macrophages infected with ΔRv2067c, Wt*Mtb* and ΔRv2067c:comp strains respectively. **c-g,** Fold enrichment of H3K9me3, H3K27me3, H3K79me3, H3K4me3 and H3K36me3 marks on peak 1 and peak 2 on the DOT1L gene in THP1 macrophages infected with *Mtb* strains. Bar pattern represents infection with following strains: Black: ΔRv2067c, Grey: Wt*Mtb*, checker:ΔRv2067c:comp. Data is represented as fold enrichment over the IgG control. **h and i,** Expression of DOT1L in HEK293T cells after transfection with scramble (lane2) and DOT1L shRNA (lane3) by western analysis and qRT-PCR. Control represents untransfected cells. β-actin was used as loading control. For qRT-PCR Ct values were normalised against GAPDH. **j,** Expression of DOT1L in uninfected THP1 macrophages (left panel), THP1 macrophages infected with *M.smeg*:FLAG (middle panel) and *M.smeg*:Rv2067c-FLAG (right panel) post treatment with EPZ004777. Inhibitor was added in increasing concentration of 0, 1.25, 2.5, 5 and 10 µM for each set. Ct values were normalised against GAPDH. All ChIP and qRT-PCR data is representative of two independent infections. For qRT PCR, two technical replicates were kept. The data plotted is mean, error bars represent s.d. and *, P-value <0.05; **,P-value <0.01; ***,P-value <0.001, ns-non significant (Student’s t-test). Values above the blot represent quantitation (arbitary units).

**Extended Data Fig. 8.**
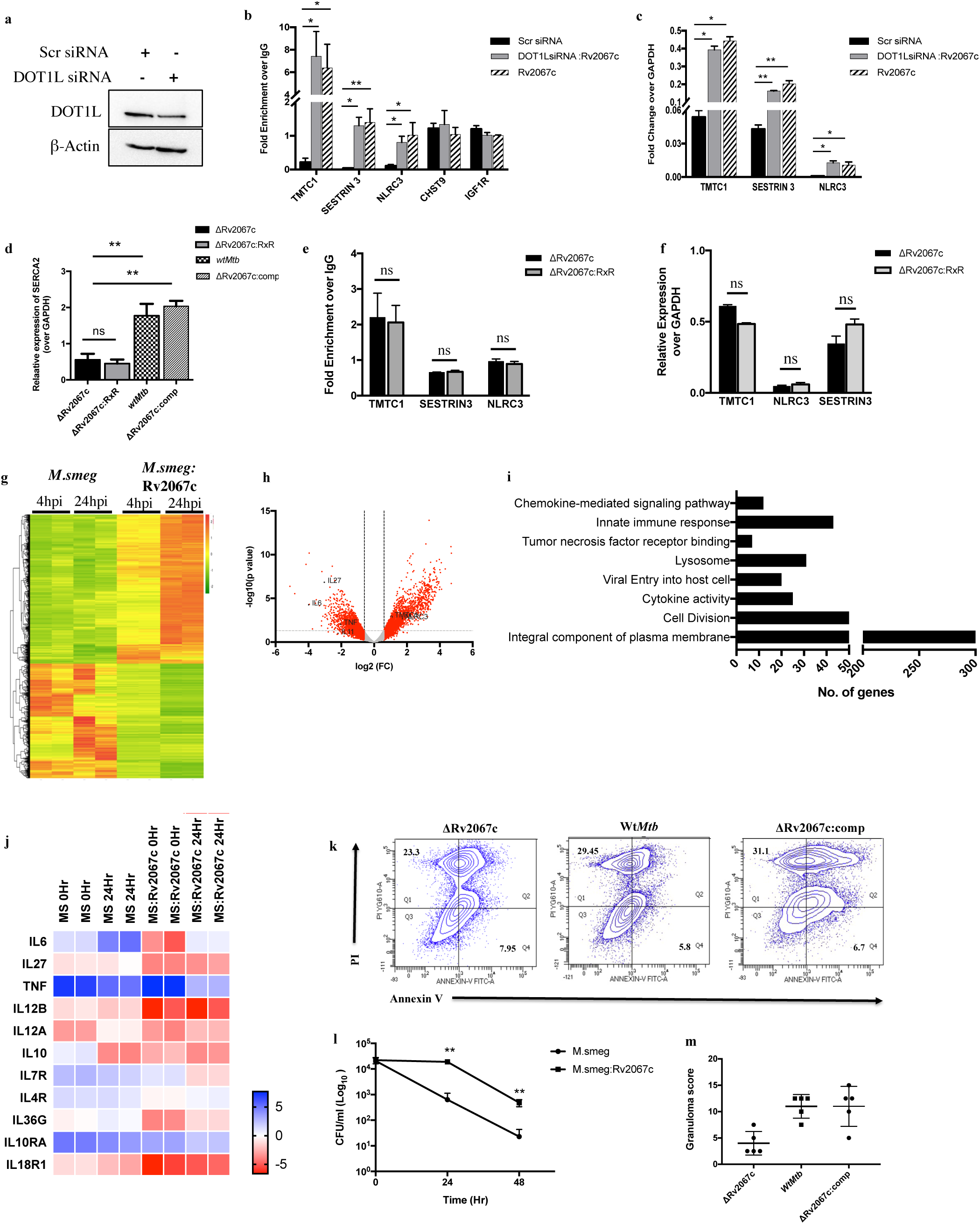
H3K79me3 mark performed by Rv2067c results in gene activation. **a,** Western blot depicts DOT1L levels in HEK293T cells treated with 20nM of scramble and DOT1L siRNA. β-Actin was kept as loading control. **b,** Bar graph depicts fold enrichment of H3K79me3 mark performed by Rv2067c on indicated gene loci in HEK-scr:pcDNA (black bar), HEK-DOT1L KD:Rv2067c (grey bar) and HEK:Rv2067c (stripe bar) cells. Data is shown as fold enrichment over the IgG control. **c,** Graph shows fold change in expression of the indicated genes in HEK-scr:pcDNA (black bar), HEK-DOT1L KD:Rv2067c (grey bar) and HEK:Rv2067c (stripe bar) cells. Ct values were normalized against GAPDH. **d,** Graph shows relative expression of SERCA2 in THP1 macrophages infected with ΔRv2067c (black bars), Wt*Mtb* (grey bars) and ΔRv2067c:comp (checker pattern) with respect to uninfected THP1 macrophages. Ct values were normalized against GAPDH. **e,** Fold enrichment of H3K79me3 mark in THP1 macrophages infected with ΔRv2067c (black bars) and ΔRv2067c:RxR (grey bars). Data is shown as fold enrichment over the IgG control. **f,** Relative expression of TMTC1, NLRC3 and SESTRIN3 in THP1 macrophages infected with ΔRv2067c (black bars) and ΔRv2067c:RxR (grey bars) with respect to uninfected THP1 macrophages. Ct values were normalized against GAPDH. **b-f,** All ChIP and qRT-PCR data is representative of two independent experiments. For qRT-PCR, two technical replicates were kept for each sample. The data plotted is mean, error bars represent s.d. and *, P-value <0.05; **,P-value <0.01; ***,P-value <0.001 (Student’s t-test). **g,** Heatmap shows relative expression profile of differentially expressed genes across the samples post infection with *M.smeg* and *M.smeg*:Rv2067c strains in macrophages. **h,** Volcano plot shows differential expression profile of genes after 24 hours of THP1 infection with *M.smeg*:Rv2067c. Red dots beyond black (vertical) and grey dotted lines (horizontal) indicate genes with fold change ≥1.5 and P-value ≤0.05. Grey dots indicate genes with fold change <1.5. Black dots show representative upregulated and downregulated genes. **i,** Bar graph shows functional annotation clusters of downregulated genes using DAVID (enrichment score >5). **j,** Heatmap depicts relative expression levels for cytokines post infection with *M.smeg* and *M.smeg*:Rv2067c strains. Log_2_RPKM values were plotted. **k,** Representative scatter plots of PI (y-axis) versus annexin V (x-axis) for macrophages infected with ΔRv2067c, Wt*Mtb* and ΔRv2067c:comp. Percentage of apoptotic and necrotic cells are indicated in Q4 and Q1 respectively. **l,** CFU analysis of THP1 macrophages infected with *M.Smeg* (circle) *and M.smeg*:Rv2067c-FLAG (square). Colony forming units were counted after 4, 24 and 48hr of infection. The data plotted is average of three independent infection. Three technical replicates were kept during each infection. Error bars represent s.d. and **,P-value <0.01 (Student’s t-test). CFU, colony forming units. **m,** Graph depicts granuloma score of lung sections of mice infected with ΔRv2067c, Wt*Mtb* and ΔRv2067c:comp, 8 weeks post infection.

**Extended Data Table 1.** Data collection and refinement statistics.

| **Dataset** | **Rv2067c**<br>**(anomalous)** | **Rv2067c-SAM**<br>**(native)** |
| --- | --- | --- |
| **Data collection** |  |  |
| Wavelength (Å) | 1.700 | 1.700 |
| Space group | P4<sub>1</sub>2<sub>1</sub>2 | P4<sub>1</sub>2<sub>1</sub>2 |
| Cell dimensions |  |  |
| a, b, c (Å) | 110.91, 110.91, 217.55 | 109.15, 109.15, 216.61 |
| α, β, γ (°) | 90, 90, 90 | 90, 90, 90 |
| Resolution | 49.41-3.25 (3.366-3.25)<sup>a</sup> | 48.74-2.40 (2.486-2.40) |
| Total reflections | 554981 (40009) | 1321981 (131545) |
| Unique reflections | 40810 (4076) | 51988 (5129) |
| R<sub>merge</sub> (%) | 0.1502 (0.9732) | 0.08841 (2.074) |
| R<sub>meas</sub> (%) | 0.156 (1.027) | 0.09024 (2.115) |
| R<sub>pim</sub> (%) | 0.04208 (0.3257) | 0.01786 (0.4132) |
| Mean I/σ(I) | 12.90 (1.8) | 25.92 (1.73) |
| CC1/2 (%) | 0.999 (0.822) | 0.999 (0.753) |
| Completeness (%) | 99.88 (99.80) | 99.96 (100) |
| Redundancy | 13.6 (9.8) | 25.4 (25.6) |
| **Refinement** |  |  |
| Resolution (Å) |  | 48.74-2.40 (2.486-2.40) |
| No. Of reflections used in refinement |  | 51978 (5129) |
| No. Of reflections used for R<sub>free</sub> |  | 2599 (257) |
| R<sub>work</sub>/R<sub>free</sub> (%) |  | 19.14/21.29 |
| No. Of non-hydrogen atoms |  | 6179 |
| Protein |  | 6120 |
| Ligand/ion |  | 10 |
| Water |  | 49 |
| Wilson B factor (Å<sup>2</sup>) |  |  |
| Average |  | 69.77 |
| Protein |  | 69.84 |
| Ligand/ion |  | 82.48 |
| Water |  | 58.72 |
| RMS deviations |  |  |
| Bond lengths (Å) |  | 0.017 |
| Bond angles (°) |  | 2.22 |
| Ramachandran plot (%) |  |  |
| Favored |  | 98.14 |
| Allowed |  | 1.86 |
| Outlier |  | 0.00 |
| Rotamer outliers (%) |  | 3.33 |
| Clash score |  | 2.81 |

Statistics for the highest-resolution shell are shown in parentheses.

**Extended Data Table 2.**
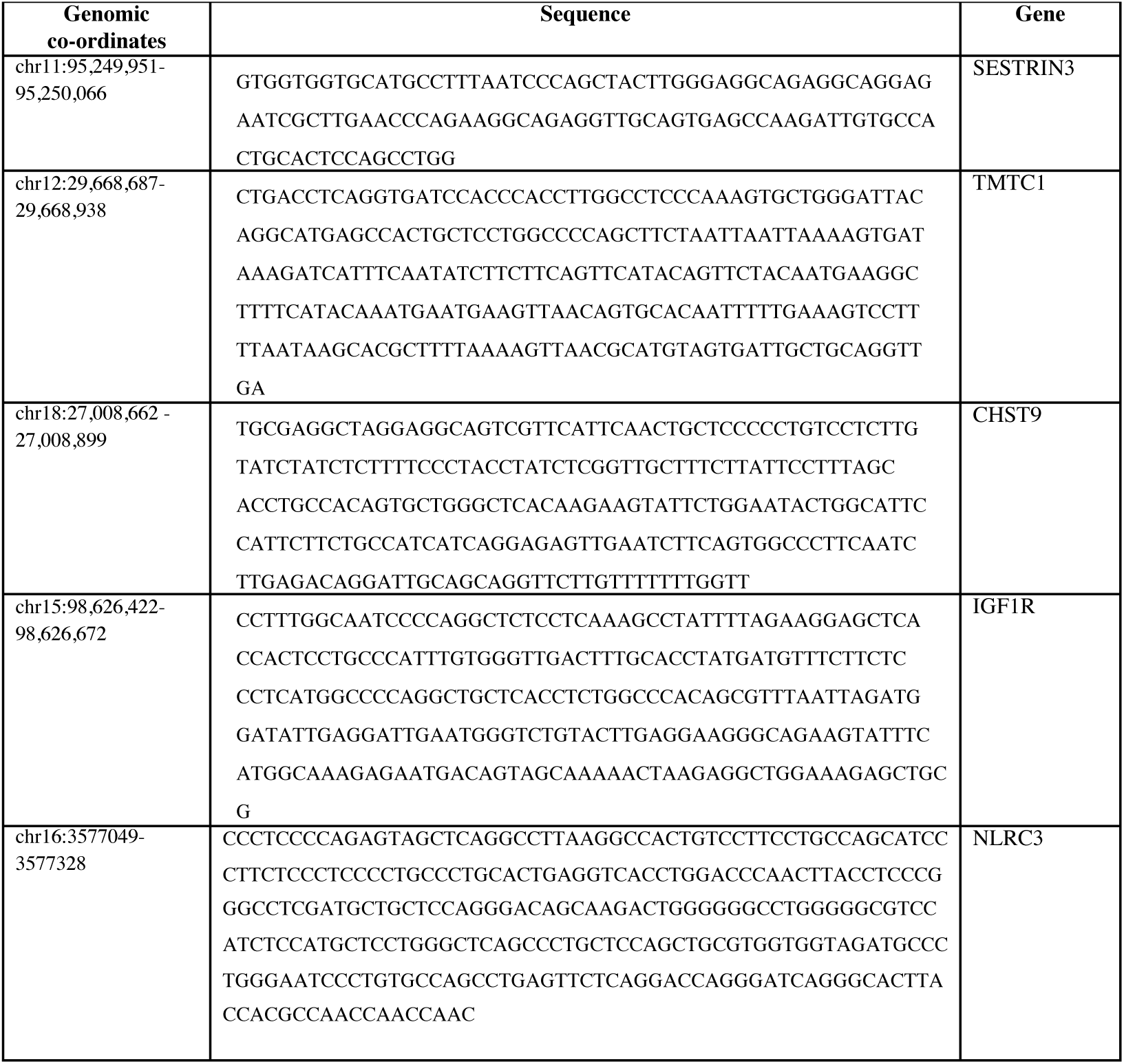
Sequence of genomic co-ordinates identified by H3K79me3 ChIP.

## Notes

### Competing Interest Statement

The authors have declared no competing interest.

