## Supplementary Results and methods for "Mycobacterium tuberculosis methyltransferase perturbs host epigenetic programming to promote bacterial survival"

**Supplementary Information**

**Results**

**Rv2067c structure determination**

The initial structure of Rv2067c was determined by experimental phasing using iodine Single wavelength Anomalous Dispersion (SAD) to 3.25 Å resolution. A better resolution structure to 2.40 Å was determined from a native dataset using the structure model obtained from experimental phasing (see Methods). The asymmetric unit (ASU) contains two molecules of Rv2067c related by a 2-fold non-crystallographic symmetry (Fig. 2a) and share an interface area of 1565.9 Å^2^ (p-value = 0.00154, calculated by PISA^1^ indicating that the crystallographic dimer is biologically relevant. The dimeric state was also confirmed by analytical gel filtration chromatography and glutaraldehyde cross linking (Extended Data Fig. 2a). The two molecules in the ASU were modeled as chain A (residues 16-208, 215-371 and 378-405) and chain B (residues 18-371 and 376-405). The remaining residues could not be modeled due to the missing electron density, an indicative of flexibility in those regions. These regions include the N-terminal segment (residues 1-15 of chia A and residues 1-17 of chain B), residues 209-214 of chain A (the corresponding residues in chain B are structured due to crystal contacts), and residues 372-377 (chain A) and residues 372-375 (chain B). An unmodeled electron density in the Fourier difference map (F_o_ – F_c_, at 3.0σ level) was observed at the putative SAM-binding site (Fig. S5). This density was modeled as S-adenosyl-L-homocysteine (SAH) due to the resulting negative density at the donor methyl carbon when modeled as SAM. However, no co-substrate (SAM or SAH) was used during purification or crystallization. Co-substrates are known to get co-purified with methyltransferases^2–4^. SAH binds in a pocket formed by a conserved motif GxG (63-GCG-65) and its interactions with protein are shown in Fig S5.

**Structural homologs of Rv2067c**

To find structures similar to Rv2067c, structural similarity search was carried out using DALI server^5^. DALI search yielded two proteins with structures similar to Rv2067c: protein lysine methyltransferase 1 from *Rickettsia prowazekii* (PKMT1, PDB: 5DO0, 5DNK and 5DPD) and protein lysine methyltransferase 2 from *R. typhi* (PKMT2, PDB: 5DOO and 5DPL) with Z-scores 27.3 and 23.2 and coverage of 342 and 345 residues, respectively^6^ . PKMT1 and PKMT2 methylate multiple lysine on outer membrane protein B (OmpB) of rickettsial species. PKMT1mainly performs monomethylation whereas PKMT2 predominantly trimethylates OmpB^7^ . The overall topology and domainal organization between Rv2067c and rickettsial PKMTs are similar except that rickettsial PKMTs contain an extra domain, middle domain, which is equivalent to the CTD of Rv2067c in terms of fold and its position in the tertiary structure (Fig. S7a,b). The putative substrate-binding cleft (termed trough, in case of Rv2067c) of PKMT1/2 is wider compared to the only Rv2067c and was characterized to be involved substrate binding^6^ .

**Methods**

**Purification of Proteins**

**Rv2067c**

Rv2067c was cloned in EcoRV digested pMyNT vector and expressed in *Mycobacterium smegmatis* mc^2^155 (*M.smeg*). *M. smeg* cells expressing Rv2067c were grown in 7H9 media (Difco) supplemented with 0.4% glucose (Sigma) and 0.05% Tween 80 (Sigma). Culture was induced at OD_600_ of 0.6 with 2% acetamide (Sigma) for 6 hr at 37°C. Cells were resuspended in lysis buffer (10 mM Tris-HCl, pH 7.4, 5% glycerol, 100 mM NaCl, 1 mM β-mercaptoethanol (BME)) containing 1 mM phenylmethylsulfonyl fluoride (PMSF) and 10 mM imidazole and lysed using French Press (SIM AMINCO) followed by short pulses of sonication (LSL SECFROID). Cell lysate was clarified by centrifuged at 13,000 ×g for 30 min and the supernatant was incubated with Ni-NTA resin (G-Biosciences). The resin was washed with lysis buffer containing 40 mM imidazole and protein was eluted with lysis buffer containing 250 mM of imidazole. Eluted fractions were checked on 12% SDS-PAGE. Pure proteins fractions were pooled, dialysed and concentrated. Rv2067c RxR mutant was cloned in pMyNT vector and purified from *M.smeg* using the above protocol.

For crystallization purpose, Rv2067c gene was subcloned into pMyNT vector to remove the cloning artifacts that arose from the vector backbone of the previous Rv2067c construct. Rv2067c gene was amplified from the previous construct using primers (Rv2067c_FP_struc and Rv2067c_RF_struc, Table S3) and assembled into linearized (with NcoI and BamHI) pMyNT vector using NEBuilder^®^ HiFi DNA Assembly mix. The new construct, pMyNT-Rv2067c-2, contains N-terminal His-tag followed by tobacco etch virus (TEV) protease cleavage site and Rv2067c open reading frame. Protein was expressed from pMyNT-Rv2067c-2 in *M. smeg* and purified as described above. The His-tag was removed using TEV protease in a cleavage buffer (50 mM Tris-HCl, pH 8.0, 50 mM NaCl, 1 mM BME) during overnight dialysis at room temperature. Cleaved tag was removed by passing the cleavage reaction through Ni-NTA column. Protein was further purified by anion exchange chromatography. Protein was loaded onto Mono-Q HiTrap (GE Healthcare) 5ml column equilibrate with low salt buffer (20 mM Tris-HCl, pH 8.0, 50 mM NaCl and 2.5 mM BME) and washed with 5 column volumes of low salt buffer. Protein was eluted with a linear gradient of 50 mM – 1000 mM NaCl. The peak containing Rv2067c fractions were pooled, concentrated and buffer exchanged to storage buffer (10mM Tris-HCl, pH 8.0, 100 mM NaCl, 2mM BME) using 10 KDa MWCO centrifugal filter (Amicon Ultra 15 mL, Millipore, USA). Protein was immediately used for crystallization trials or snap frozen in liquid N_2_ and stored at -80ºC for future use.

**Histone H3**

Histone H3 cloned in NdeI–HindIII restriction sites of pET28b (+) vector was purified from *E. coli* BL21 (DE3) using 0.3mM IPTG induction for 4 hr at 37°C. Cells were resuspended in lysis buffer (50mM Tris pH 7.4, 250mM NaCl and 0.1% sarcosine). Lysed cells were centrifuged at 13,000 rpm for 30 min, the supernatant was incubated with Ni-NTA beads. The protein was eluted in elution buffer containing 350 mM of imidazole. Pure proteins fractions were pooled, dialysed and concentrated.

**Human core histones (H3, H4, H2A and H2B)**

Human core histone proteins H3, H4, H2A and H2B were expressed from the constructs pET21a-H3, pET3a-H4, pET21a-H2A and pET21a-H2B, respectively. *E. coli* BL21(DE3) Rosetta2 cells harboring the above plasmids were grown in 2×YT broth containing 0.1% w/v glucose, ampicillin (100 µg/mL) and chloramphenicol (25 µg/mL) at 37°C. Cells were induced at OD_600_ between 0.6 and 0.8 for 2 hr (H3 and H4) or 3 hr (H2A and H2B) with 0.3 mM IPTG. Cells were harvested by centrifugation at 6,000×g and resuspended in a resuspension buffer (50 mM Tris-HCl, pH 7.4, 100 mM NaCl, 1 mM PMSF and 2.5 mM BME). Cell lysis was carried out by alternate freeze-thaw cycles followed by sonication. Lysate was centrifuged at 23,000×g at 4°C for 20min. Pellet containing inclusion bodies was washed twice with wash buffer (50 mM Tris-HCl, pH 7.4, 100 mM NaCl, 1 mM EDTA, 1 mM PMSF and 2.5 mM BME) containing 1% v/v Triton X-100 followed by two washes without TritonX-100. Histones were purified from these washed inclusion bodies by acid-extraction^8^. In brief, inclusion bodies were homogenized in 10 mL of 0.25 N HCl and incubated at -20ºC for 30min. The insoluble portion was removed by centrifugation at 27,000×g, 4ºC for 10 min and the supernatant containing histones was neutralized with 0.125 volumes of 2 M Tris. All the histones were dialysed overnight against water containing 2 mM BME at 4ºC using 6,000 KDa MWCO dialysis membrane. The dialysed histones were lyophilized and stored at -20ºC for future use. For MTase assays, these histones were further purified by gel filtration followed by cation-exchange chromatography as described^9^.

**DOT1L^2-416^**

Human DOT1L construct (pEW3:pET32a DOT1L(2-416)) was purchased from Addgene, USA (Plasmid#124098, https://www.addgene.org/124098/). Expression and purification of DOT1L protein (DOT1L^2-416^) was carried out as described^10^.

**Preparation of Widom 601 DNA**

pUC57-Widom 601 plasmid containing eight repeats of Widom 601 sequence^11^, a 145 bp strong nucleosomal positioning DNA sequence, flanked by EcoRV sites was produced in *E. coli* Top10 cells. Plasmid isolation, EcoRV digestion and purification of Widom 601 sequence were carried out as described^12^.

**Preparation of nucleosome core particles (NCPs)**

Histone octamers were refolded from core histones using salt dialysis method as described ^13^. Since all the four histone preparations contained impurities, the estimation of concentrations was not accurate. Thus, the apparent quantities of H2A and H2B were kept 1.5 molar excess to H3 and H4 during reconstitution. This eases the purification of octamers from histone tetramers and hexamers by size exclusion chromatography^14^. All four histones were dissolved in unfolding buffer (20 mM Tris-HCl pH, 7.5, 6 M guanidinium chloride, 5 mM DTT) and dialysed three times against refolding buffer (10 mM Tris-HCl, pH 7.5, 2 M NaCl, 1 mM Na-EDTA, 5 mM BME) at 4ºC. The precipitated protein was removed by centrifugation at 22,000×g, 4ºC for 30 min. The supernatant was concentrated and loaded onto gel filtration column (Superdex 200) preequilibrated with refolding buffer, at 0.8 ml/min flow rate. Fractions were checked on SDS-PAGE (Extended Data Fig. 1g). Fractions with equimolar concentration of all four histone were pooled, concentration was determined and stored at -20ºC with 50% glycerol.

NCPs were reconstituted from octamers and Widom 601 DNA using microscale reconstitution protocol^12^. For reconstitution, 1.66 µg of DNA was mixed with 2.4 µg of octamers (molar ratio of DNA to octamers 1:1.2) to a final 2 M NaCl concentration in 10 µl volume and incubated on ice for 30 min. The salt concentration was slowly reduced to 100 mM in the reconstitution mix by sequential addition of 10 µl, 5 µl, 5 µl, 70 µl and 100 µl of 10 mM Tris-HCl, pH 7.6 with 1 hr incubation on ice at every step. Multiple such microscale preparations were pooled and concentrated using centrifugal filter. The quality of the preparation was assessed on 6% native PAGE (Extended Data Fig. 1h).

**Sequence analysis**

Homologous protein sequences of Rv2067c were obtained by BLAST^15^ against NCBI Reference Protein (RefSeq_Protein) database. BLAST hits with more than 80% sequence coverage resulted in minimum sequence identity of ∼ 23% were used for analysis. Two sequence sets (A and B) were generated from these hits. Set A contains sequences only from mycobacterial species clustered at 80% identity and set B contains all hits clustered at 80% identity. CD-HIT^16^ was used for clustering. Multiple sequence alignment (MSA) for both the sets was generated using Clustal Omega^17^. Residue-wise conservation scores were calculated from MSA using ConSurf^18^ with default parameters and the conservation scores were mapped onto Rv2067c structure. A few selected sequences from set B were used to generate representative MSA and was rendered using Jalview^19^.

**Molecular dynamics simulations**

Crystal structure of Rv2067c dimer with cofactor, SAM, was used for simulations. Prior to simulations, missing loops were built and SAH was replaced with SAM using Coot^20^. The missing N-terminus residues (chain A: residues 1-15 and chain B: residues 1-17) were modeled by grafting these residues from Rv2067c model generated using AlphaFold2 (AF2)^21^. Thus modeled Rv2067c-SAM dimer was solvated with TIP3P water model^22^ in a dodecahedron box with 1.2 nm padding from the protein atoms. The charge of the system was neutralized while keeping the concentration of Na^+^ and Cl^–^ ions at 0.15 M. Simulation system was parameterized using CHARMM36m force field^23^ which also contains force field parameters for SAM. Energy minimization (50,000 steps) was carried out with steepest-descent method followed by equilibration in NVT and NPT ensembles, 100 ps each, with positional restraints (1000 kJ·mol^-1^·nm^-2^). Unrestrained production simulations of 100 ns were carried out in NPT ensemble at 300 K temperature and 1 bar pressure. Temperature was maintained by velocity rescaling scheme^24^ and pressure with Parrinello-Rahman barostat^25^ . Neighbor search was carried out using Verlet cutoff-scheme with neighbor list updated every 40 steps and van der Waals interactions were calculated up to 1.2 nm radius. Long range electrostatics (1.2 nm cut-off distance) were computed using Particle Mesh Ewald method with cubic interpolation and 1.6 nm Fourier spacing. Bonds to hydrogen atoms were constrained using LINCS algorithm. Two femtosecond time steps were used for integration. Simulation trajectory was saved at 2 ps time interval. Simulations were carried out using GPU-accelerated Gromacs 2021.2^26^. The trajectory was processed using tools in Gromacs package and CPPTRAJ^27^ from Amber package ([http://ambermd.org](http://ambermd.org/)).

**Analysis of putative substrate-binding trough**

MD trajectory was corrected for periodic boundary condition. Each monomer of the dimer was written to a separate trajectory, at 10 ps time intervals, without water molecules and ions. Frames of the trajectory were aligned to the crystal structure using three different sets of backbone atoms (CA, C, N, O, and H) viz. all residues (407-aa), seven-β-strand core (7BS-aa) and 7BB-aa and part of LSD (7BS + LSD). The aligned set with minimum RMSF (root mean square fluctuations) around the putative active site region i.e., 7BS-aa + LSD was chosen (Fig. S3) for calculation of volumetric density map of the putative substrate-binding trough using POVME 3.0^28^. A custom inclusion volume of the grid with 1.0 Å spacing encompassing the putative substrate-binding trough was defined for volume measurement in POVME 3.0. Another custom inclusion volume around the putative active site pocket was defined to estimate the widened active site volume. As a crude estimate, frames with volume more than or equal to the side chain volume of lysine were considered as widened active site and used for preparation of simulation movie using ChimeraX^29^ .

**Enzyme-substrate reaction complex model, rotation-scan, and rationale**

A model of reaction-complex for an enzymatic reaction was constructed with respect to protein lysine methylation. This model depicts the complex between the enzyme and its substrate in an imminent enzymatic reaction, i.e., when the reaction is about to take place. The following assumptions were made for constructing the enzyme-substrate reaction-complex model. During any enzymatic reaction, an enzyme-substrate complex is formed without steric hindrance between the enzyme and its substrate and is pre-deterministic for a given enzyme-substrate complex. In an imminent reaction, the reacting atoms from both the enzyme and its substrate must come in contact for a reaction to take place. Protein lysine methylation (a SAM-dependent methylation) follows a bimolecular nucleophilic substitution (SN_2_) reaction mechanism where methyl acceptor atom of the substrate, the ζ-nitrogen (NZ^SUB^ ), attacks methyl carbon of SAM (CE^SAM^ ), nucleophilically. In an imminent reaction, NZ^SUB^ lies on an axis that passes through the scissile bond, the bond between sulphur (SD^SAM^ ) and CE^SAM^ atoms, and in contact with CE^SAM^ at a van der Waals (vdW) contact distance 3.30 Å^30–32^ (Fig. S7). Here, we termed the position of NZ SUB atom as the ‘reaction enter’ of a reaction-complex. All the possible orientations of the substrate with respect to its enzyme, in a reaction-complex, were determined with the help of Euler’s rotation theorem ^33^. The substrate was rotated about the reaction center, while the enzyme was fixed, using a rotation matrix (R), given by eq. 1. (1)

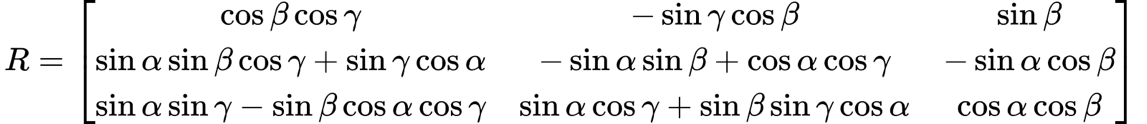
(1)

The α, β and γ are the Tait-Bryan angles of elemental rotations about x, y, and z, respectively. A range of angles α = -180 - +180, β = -90 - +90 and γ = -180 - +180, that constitutes a rotation-scan, is sufficient to obtain all the possible orientations of the substrate with respect to the enzyme. Prior to the rotation-scan, the reaction center was shifted to the origin to make it rotation invariant, about which either substrate or enzyme is rotated. All the transformations were carried out in a Cartesian coordinate system. For each rotation (a combination of α, β and γ), the total number of clashing atoms (TCA) between the enzyme and its substrate were calculated using the criterion given by equation 2^32^.

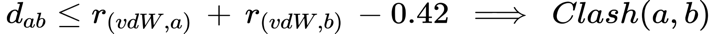
  (2)

 The *d_ab_* is the distance between the atom *a* of the substrate and the atom *b* of the enzyme, and *r_(vdW,a)_* and *r_(vdW,b)_* are the vdW radii of atoms *a* and *b*, respectively. Atom pairs (*ab*) that follow the clash criterion were added up to TCA. The TCA was used as a metric to choose the orientation with minimal clashes. For simplicity, all hydrogen atoms were excluded from the calculations. The vdW radii for atoms were chosen from Word, J. M. et al^32^. Carbonyl carbon vdW radius was set as same as vdW radius of non-carbonyl carbon. Rotation-scan was implemented using python script.

**Construction of nucleosome-Rv2067c and nucleosome-DOT1L reaction-complex models for rotation-scan**

Rv2067c-nucleosome complex model was constructed by imposing the assumptions described for enzyme-substrate reaction complex model. Coordinates for Rv2067c and nucleosome were taken from the crystal structure of Rv2067c and cryo-electron microscopy structure of UbNuc-DOT1L active-state complex (PDB: 6NJ9), respectively. In Rv2067c structure, SAM was modeled in place of SAH using Coot^20^. A lysine analog, norleucine (Nle), at H3K79 position (H3K79Nle) in 6NJ9 was mutated, in silico, to an extended rotamer of lysine. The reaction-center was defined within Rv2067c. Coordinates of the Rv2067c and nucleosome were transformed such that the reaction-center and the NZ atom of the H3K79 (NZ^H3K79^) of nucleosome were at the origin. The initial orientations of Rv2067c and nucleosome, in the Nucleosome-Rv2067ccomplex model, were set by transforming the centroids of Rv2067c and nucleosome onto the positive and negative y-axis, respectively. The active-state complex of UbNuc-DOT1L (PDB: 6NJ9) was used for benchmarking the rotation-scan method. To construct the nucleosome-DOT1L reaction complex, all ubiquitin and one DOT1L molecule (chain M) were removed from 6NJ9. H3K79Nle was mutated to lysine, in silico. The position of NZ^H3K79^ (methyl acceptor) was treated as reaction-center and was moved to the origin. Three different starting models of Nuc-DOT1L with randomly oriented DOT1L, that are different from the experimental complex (6NJ9), were generated. All the models were subjected to rotation-scan. The TCA were plotted as a function of any two elemental rotation angles (α or β or γ) while the third angle denotes the minimum TCA value. In a benchmarking test, all the three randomly oriented DOT1L molecules could generate DOT1L binding conformation that is seen in nucleososme-DOT1L complex, with the least number of clashing atoms, in a rotation-scan (Fig. S8)

**Modeling of Rv2067c - H3 peptide (73-83) complex**

H3 peptide (73-83) was manually modeled, using Coot^20^, into the putative substrate-binding trough of Rv2067c in two binding modes, based on the criteria followed for the reaction center and the methyl acceptor residue binding to the protein methyltransferases. The substrate lysine (H3K79) was placed nearly perpendicular to the long axis of SAM with a spatial constraint while the NZ^H3K79^ atom lies at the reaction center. The complete peptide was built by the addition of amino acid residues to the N- and C-termini of thus placed lysine. In one mode peptide orients in the direction of N- to C-termini whereas in the other it is C- to N-termini, along the trough. Two peptides with one mode per monomer of Rv2067c dimer were built. Thus, built Rv2067c-H3 peptide complex was energy minimized, in vacuum, using pmemd.cuda module of Amber package^34^. The system was parameterized using amber FF14SBforce field^35^. Force filed parameters for SAM were obtained from Saez D et al ^36^.

**Software**

Structures were visualized using ChimeraX^29^, plotting was done using Matplotlib, and figures schematics were made using Inkscape.

**Glutaraldehyde cross linking and analytical gel filtration**

For crosslinking, Rv2067c was treated with various concentrations of glutaraldehyde in buffer containing 20 mM HEPES, pH 7.5 and 50mM KCl and incubated at room temperature for 15 min. Reactions were terminated with 1X SDS dye and analysed on 8% SDS PAGE, with detection by [silver staining](https://www.sciencedirect.com/topics/biochemistry-genetics-and-molecular-biology/silver-staining).

For analytical gel filtration, Rv2067c was loaded on to a 24 ml Superdex 200 increase column (GE Healthcare). The standard curve was prepared from the retention volumes of ribonuclease A (13.7kDa), Ovalbumin (43kDa), Conalbumin (75kDa), Aldolase (158kDa), Ferritin (440kDa) and used for determining the molecular weight of Rv2067c from its retention volume by linear interpolation.

**RNA and genomic DNA isolation from Mycobacteria**

For RNA and genomic DNA isolation, *Mtb* strains were grown to an OD_600_ of 0.6-0.8. Cells were harvested and washed once with PBS. RNA extraction was conducted using the FastRNA® Pro Blue Kit (MP Biomedicals, USA) in accordance with the manufacturer's instruction. For genomic DNA isolation, bacteria was heat killed and DNA was extracted using CTAB, followed by phenol extraction and ethanol precipitation^37^.

**Annexin V and Propidium Iodide (PI) Staining**

THP1 macrophages seeded at a density of 0.5 x10^6^ were infected with Wt*Mtb,* ΔRv2067c and ΔRv2067c:comp. Cell death by apoptosis or necrosis in macrophages infected with different strains was quantified using Dead Cell Apoptosis Kits with Annexin V for Flow Cytometry (Invitrogen). Apoptosis was measured as Annexin V-FITC single positive (Q4) and necrosis as PI single positive (Q1). The flow cytometry assays were performed using FACSAria Fusion (BD Biosciences) and the data were analysed using FACSDiva.

**RNA sequencing analysis**

Data quality was checked using FastQC^38^ and MultiQC^39^ software. The data was checked for base call quality distribution. Raw sequence reads were processed to remove adapter sequences and low-quality bases using fastp^38^. The QC passed reads were mapped onto indexed Human reference genome (GRCh38.p7) using STAR2^40^ aligner. Gene level expression values were obtained as read counts using featureCounts software^41^ .

Multistep analysis was performed to identify differential regulated genes by Rv2067c. First, differentially expressed gene-set was identified for macrophages infected with *M.smeg* expressing Rv2067c at 24 h.p.i in comparison to 4 h.p.i. Similarly, a gene-set was identified for macrophages infected with *M.smeg*. Next, the two gene sets were compared to identify deregulated genes by Rv2067c post 24 h.p.i. Differential expression analysis was carried out using edgeR^42^ package after normalizing the data based on trimmed mean of M (TMM) values.
