## Supplementary Figures 1-8 for "Mycobacterium tuberculosis methyltransferase perturbs host epigenetic programming to promote bacterial survival"

### Slide 1
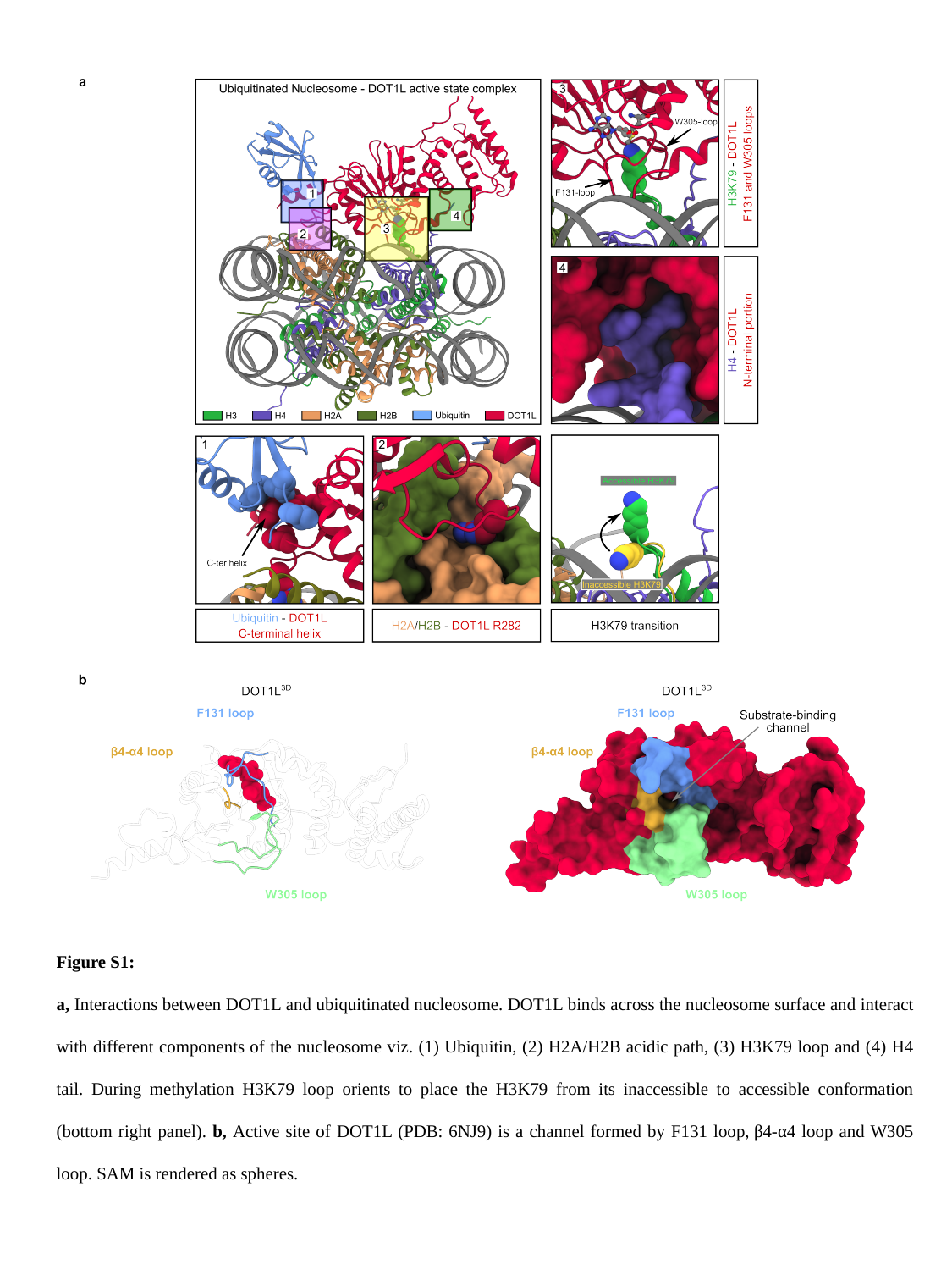

Figure S1:
a, Interactions between DOT1L and ubiquitinated nucleosome. DOT1L binds across the nucleosome surface and interact with different components of the nucleosome viz. (1) Ubiquitin, (2) H2A/H2B acidic path, (3) H3K79 loop and (4) H4 tail. During methylation H3K79 loop orients to place the H3K79 from its inaccessible to accessible conformation (bottom right panel). b, Active site of DOT1L (PDB: 6NJ9) is a channel formed by F131 loop, β4-α4 loop and W305 loop. SAM is rendered as spheres.

### Slide 2
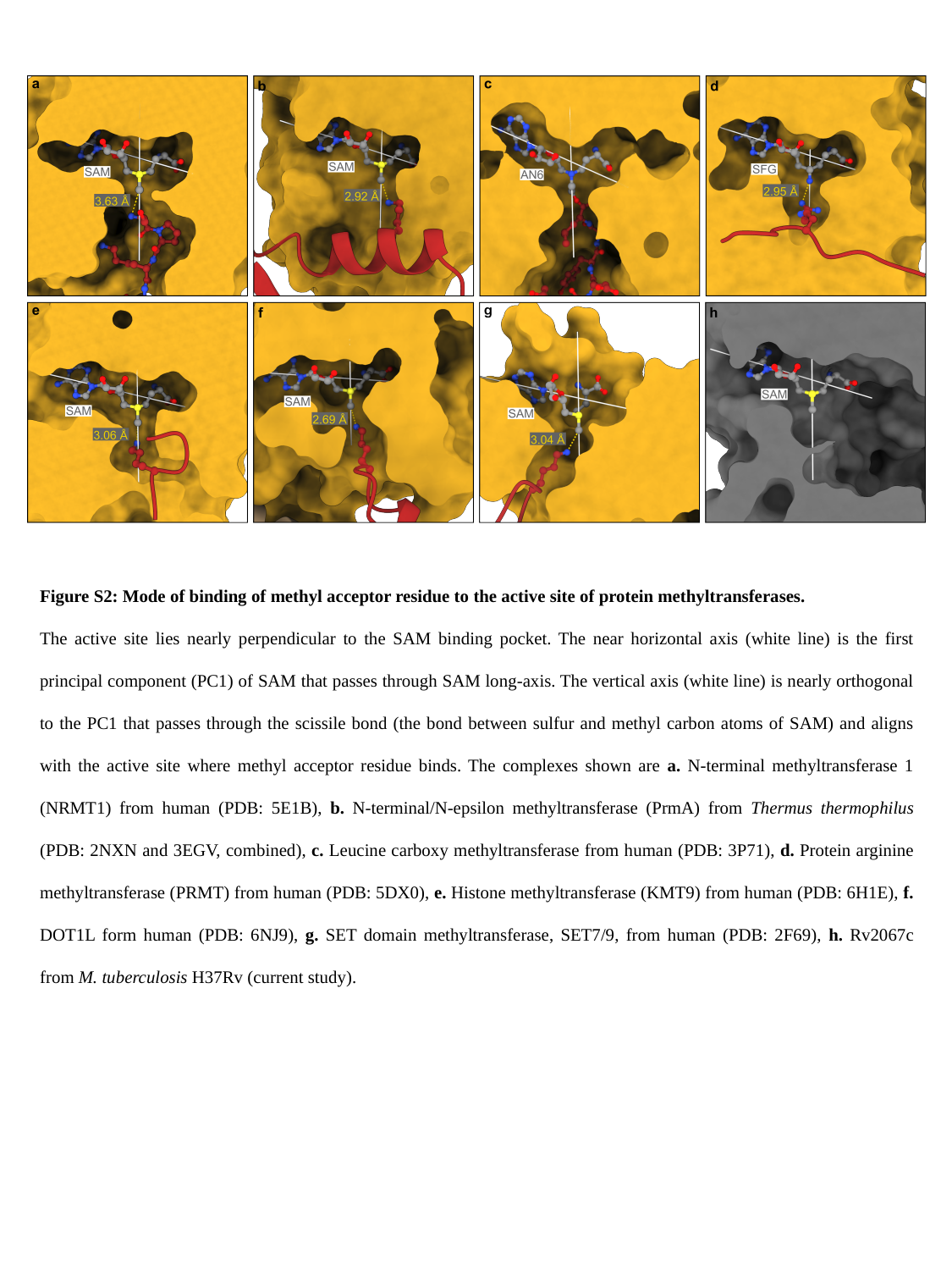

Figure S2: Mode of binding of methyl acceptor residue to the active site of protein methyltransferases.
The active site lies nearly perpendicular to the SAM binding pocket. The near horizontal axis (white line) is the first principal component (PC1) of SAM that passes through SAM long-axis. The vertical axis (white line) is nearly orthogonal to the PC1 that passes through the scissile bond (the bond between sulfur and methyl carbon atoms of SAM) and aligns with the active site where methyl acceptor residue binds. The complexes shown are a. N-terminal methyltransferase 1 (NRMT1) from human (PDB: 5E1B), b. N-terminal/N-epsilon methyltransferase (PrmA) from Thermus thermophilus (PDB: 2NXN and 3EGV, combined), c. Leucine carboxy methyltransferase from human (PDB: 3P71), d. Protein arginine methyltransferase (PRMT) from human (PDB: 5DX0), e. Histone methyltransferase (KMT9) from human (PDB: 6H1E), f. DOT1L form human (PDB: 6NJ9), g. SET domain methyltransferase, SET7/9, from human (PDB: 2F69), h. Rv2067c from M. tuberculosis H37Rv (current study).

### Slide 3
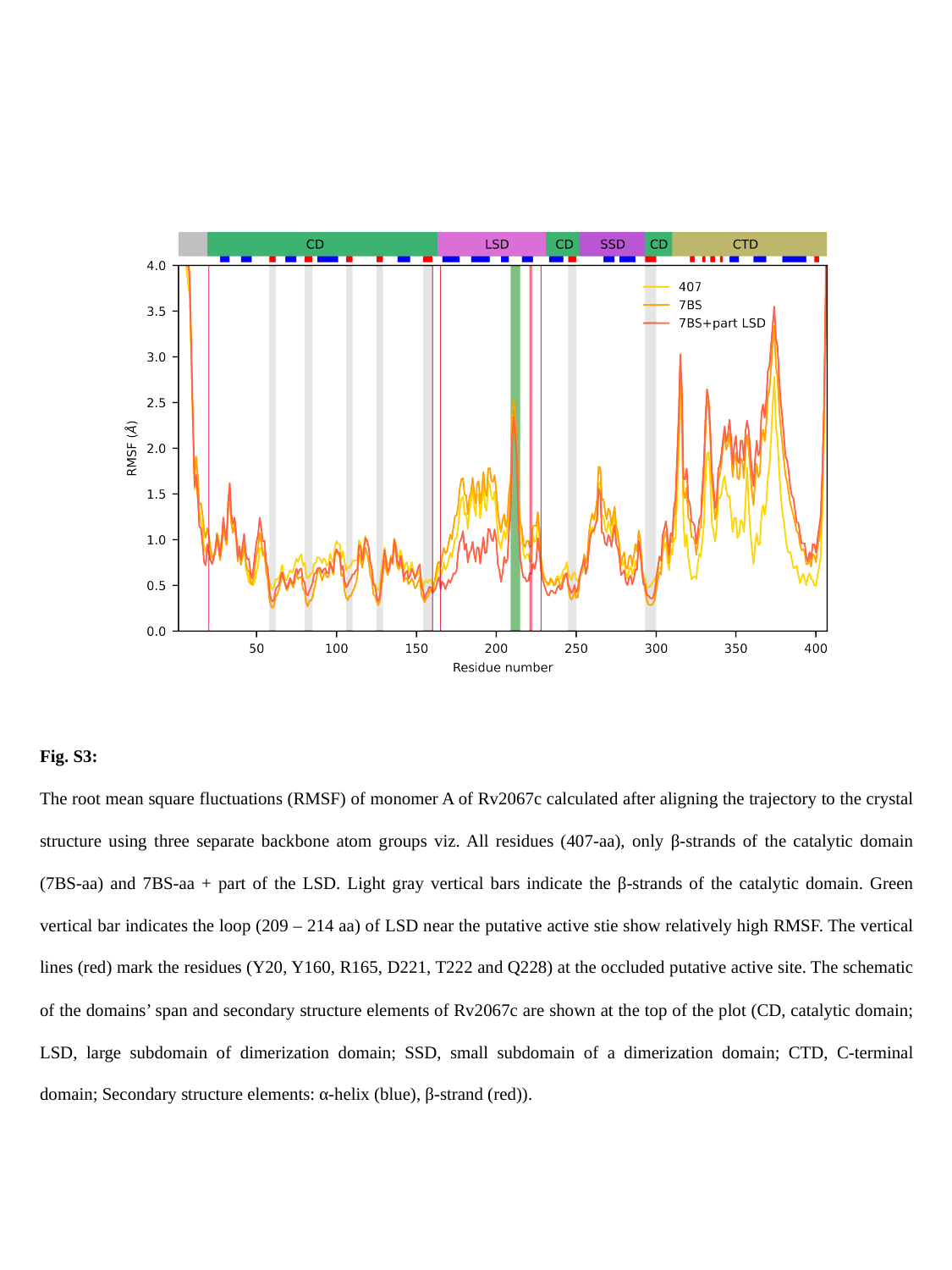

Fig. S3:
The root mean square fluctuations (RMSF) of monomer A of Rv2067c calculated after aligning the trajectory to the crystal structure using three separate backbone atom groups viz. All residues (407-aa), only β-strands of the catalytic domain (7BS-aa) and 7BS-aa + part of the LSD. Light gray vertical bars indicate the β-strands of the catalytic domain. Green vertical bar indicates the loop (209 – 214 aa) of LSD near the putative active stie show relatively high RMSF. The vertical lines (red) mark the residues (Y20, Y160, R165, D221, T222 and Q228) at the occluded putative active site. The schematic of the domains’ span and secondary structure elements of Rv2067c are shown at the top of the plot (CD, catalytic domain; LSD, large subdomain of dimerization domain; SSD, small subdomain of a dimerization domain; CTD, C-terminal domain; Secondary structure elements: α-helix (blue), β-strand (red)).

### Slide 4
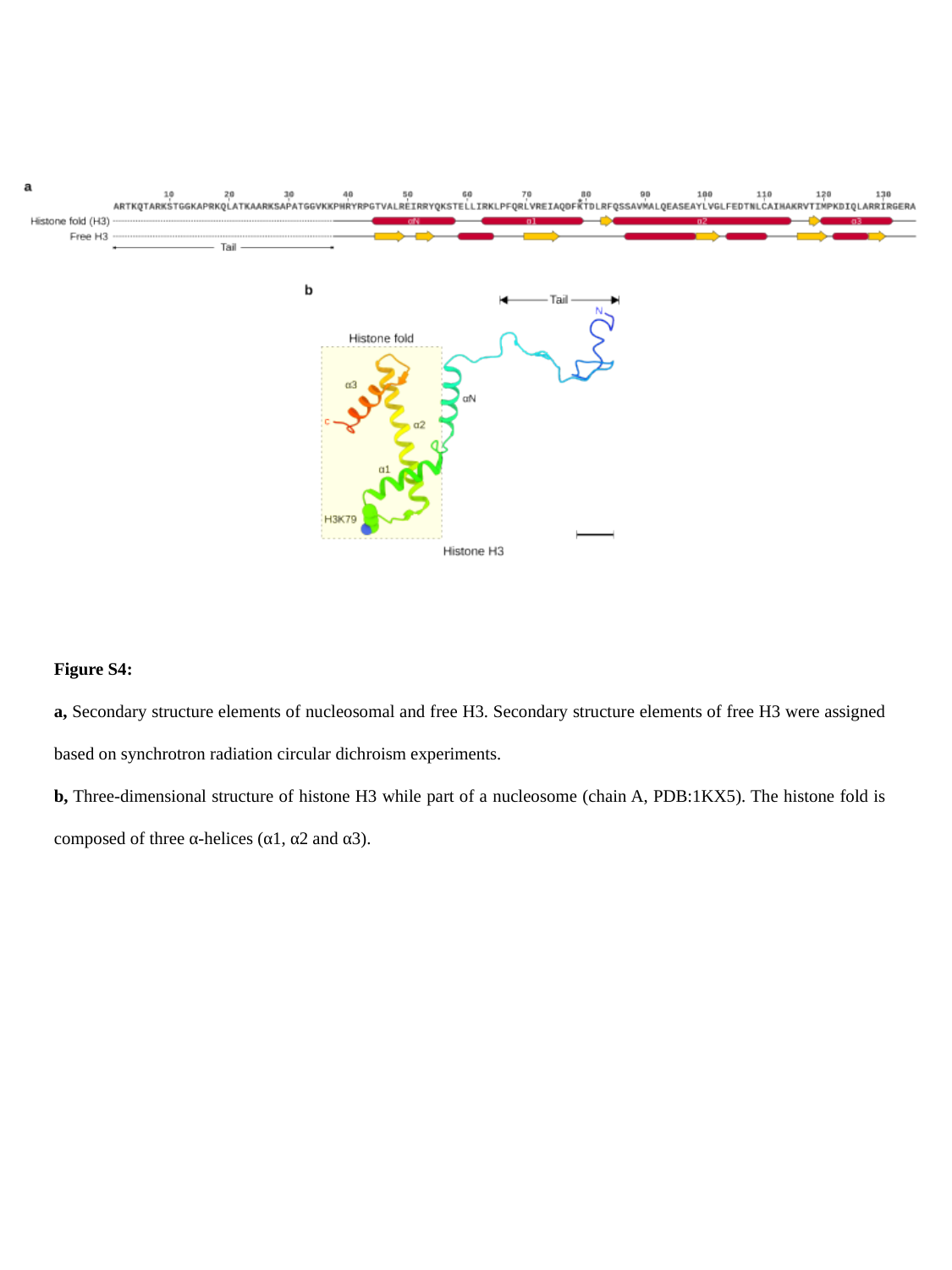

Figure S4:
a, Secondary structure elements of nucleosomal and free H3. Secondary structure elements of free H3 were assigned based on synchrotron radiation circular dichroism experiments.
b, Three-dimensional structure of histone H3 while part of a nucleosome (chain A, PDB:1KX5). The histone fold is composed of three α-helices (α1, α2 and α3).

### Slide 5
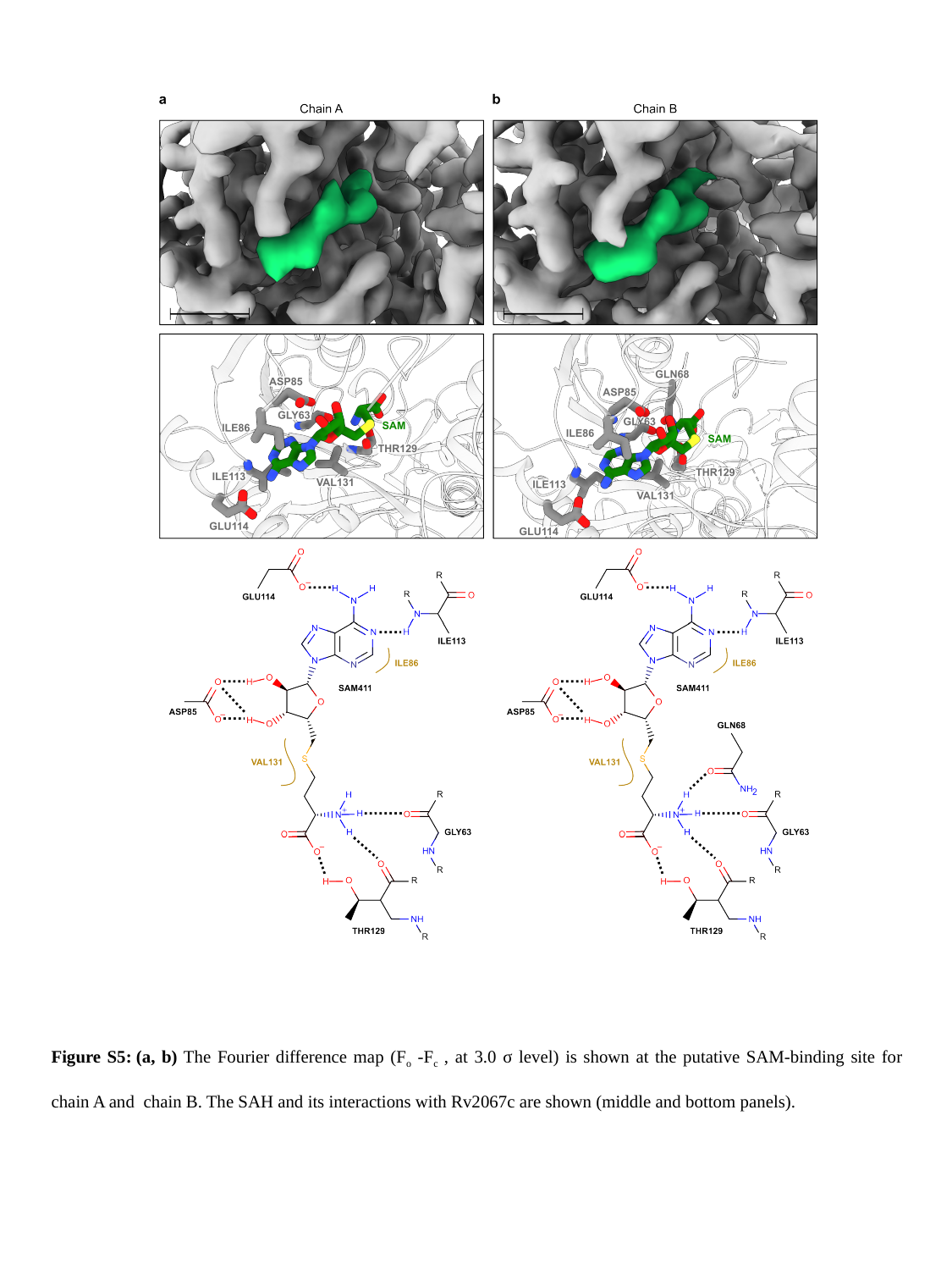

Figure S5: (a, b) The Fourier difference map (F_o_ -F_c_ , at 3.0 σ level) is shown at the putative SAM-binding site for chain A and  chain B. The SAH and its interactions with Rv2067c are shown (middle and bottom panels).

### Slide 6
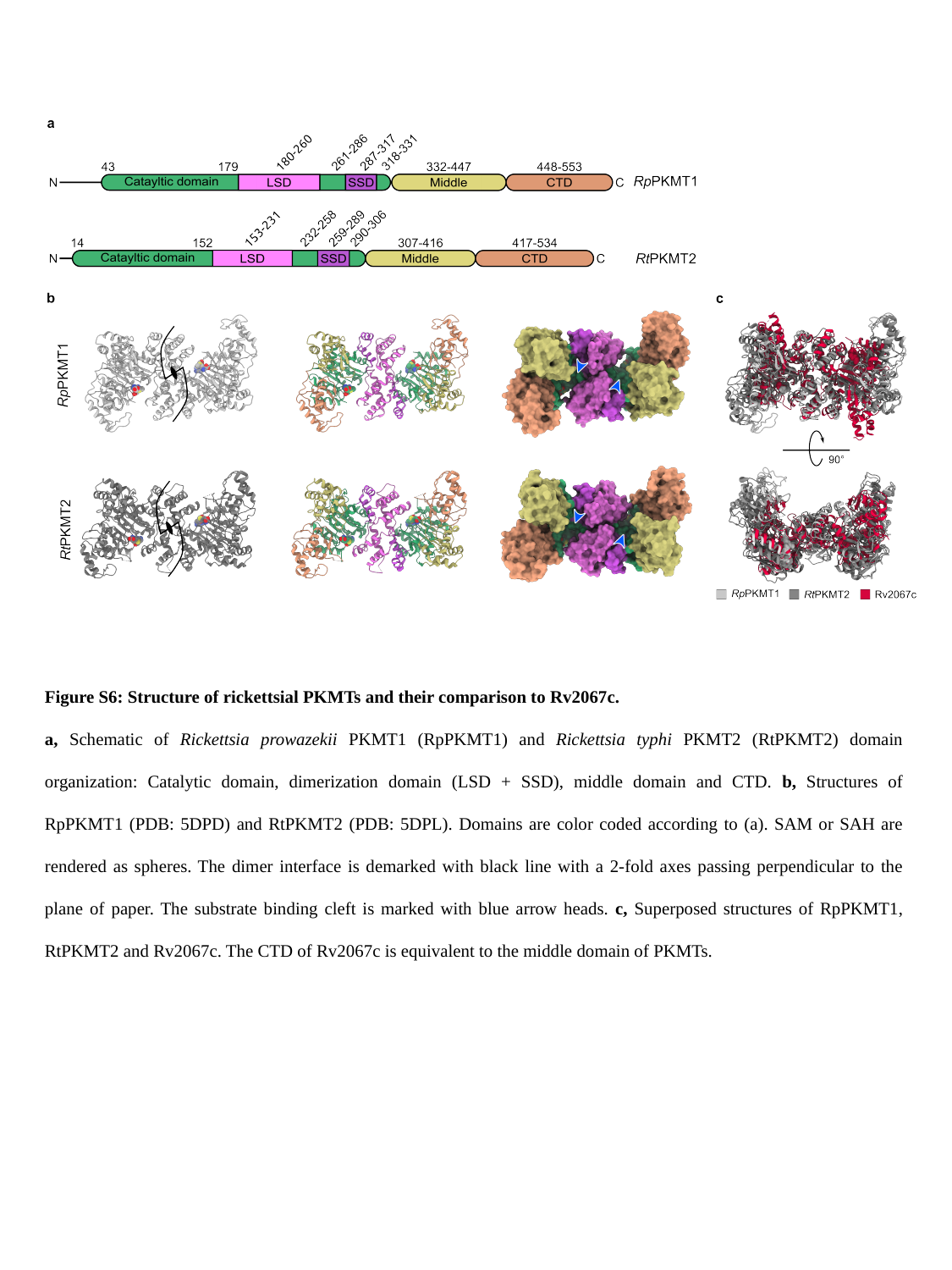

Figure S6: Structure of rickettsial PKMTs and their comparison to Rv2067c.
a, Schematic of Rickettsia prowazekii PKMT1 (RpPKMT1) and Rickettsia typhi PKMT2 (RtPKMT2) domain organization: Catalytic domain, dimerization domain (LSD + SSD), middle domain and CTD. b, Structures of RpPKMT1 (PDB: 5DPD) and RtPKMT2 (PDB: 5DPL). Domains are color coded according to (a). SAM or SAH are rendered as spheres. The dimer interface is demarked with black line with a 2-fold axes passing perpendicular to the plane of paper. The substrate binding cleft is marked with blue arrow heads. c, Superposed structures of RpPKMT1, RtPKMT2 and Rv2067c. The CTD of Rv2067c is equivalent to the middle domain of PKMTs.

### Slide 7
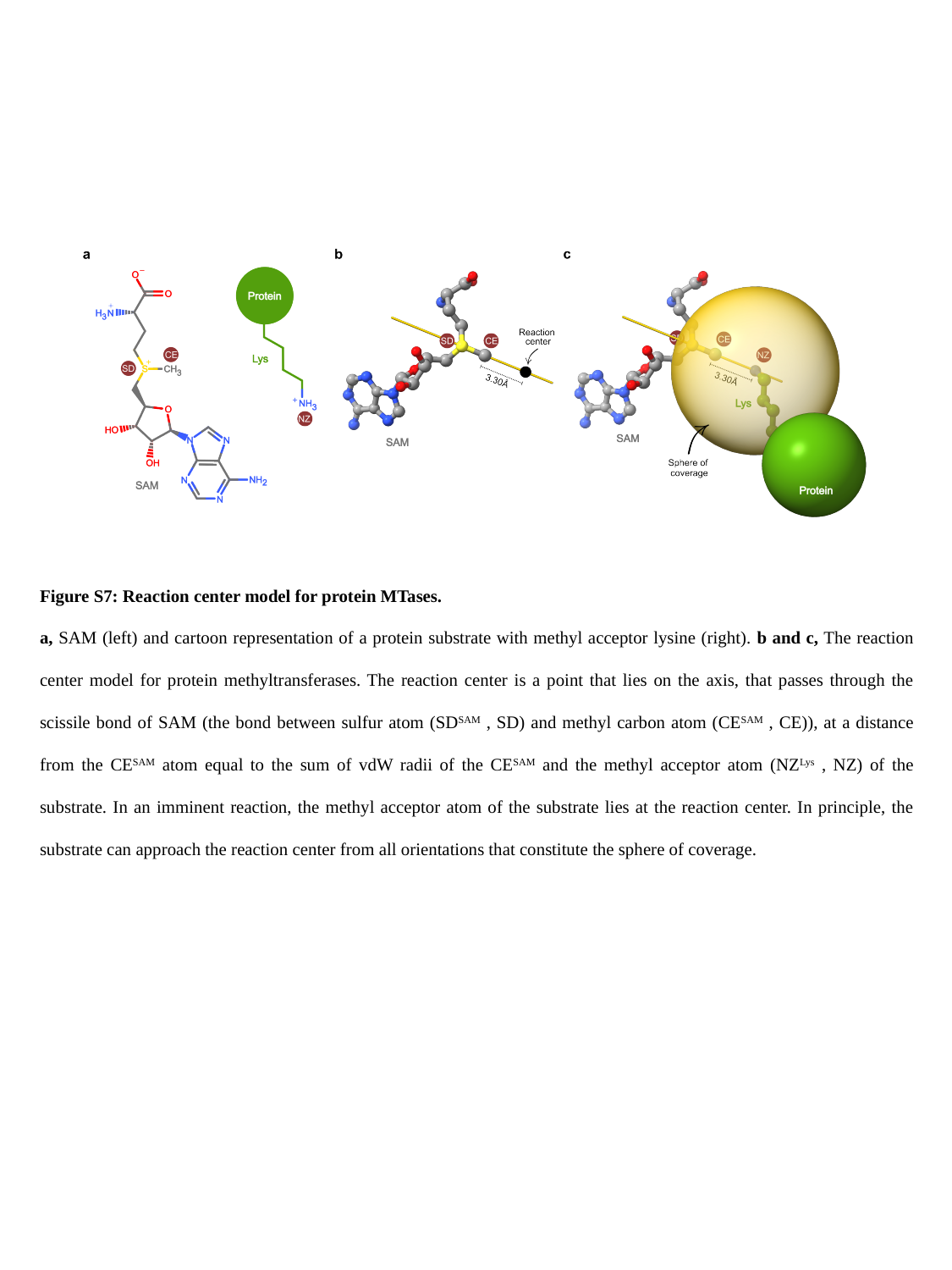

Figure S7: Reaction center model for protein MTases.
a, SAM (left) and cartoon representation of a protein substrate with methyl acceptor lysine (right). b and c, The reaction center model for protein methyltransferases. The reaction center is a point that lies on the axis, that passes through the scissile bond of SAM (the bond between sulfur atom (SD^SAM^ , SD) and methyl carbon atom (CE^SAM^ , CE)), at a distance from the CE^SAM^ atom equal to the sum of vdW radii of the CE^SAM^ and the methyl acceptor atom (NZ^Lys^ , NZ) of the substrate. In an imminent reaction, the methyl acceptor atom of the substrate lies at the reaction center. In principle, the substrate can approach the reaction center from all orientations that constitute the sphere of coverage.

### Slide 8
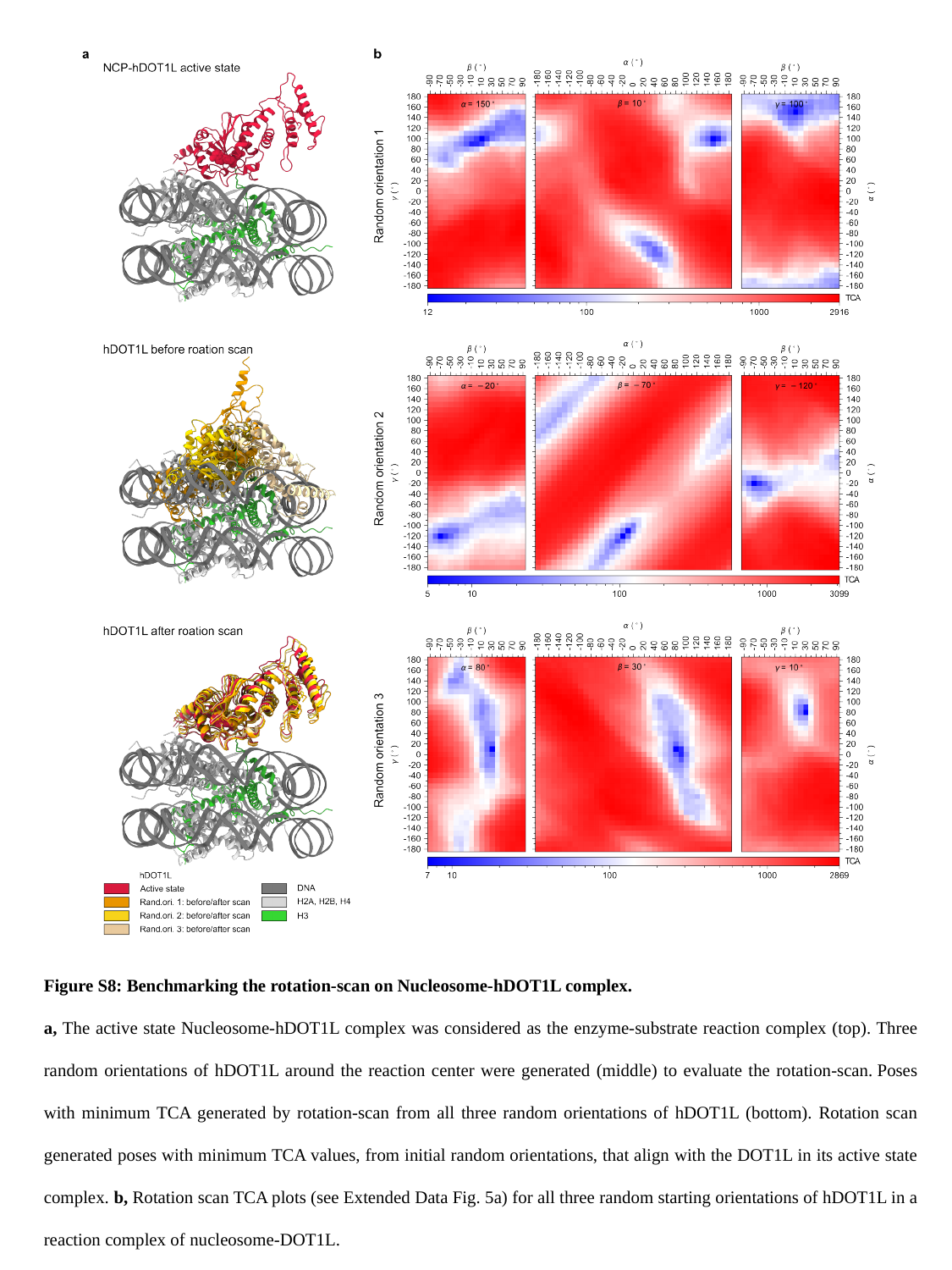

Figure S8: Benchmarking the rotation-scan on Nucleosome-hDOT1L complex.
a, The active state Nucleosome-hDOT1L complex was considered as the enzyme-substrate reaction complex (top). Three random orientations of hDOT1L around the reaction center were generated (middle) to evaluate the rotation-scan. Poses with minimum TCA generated by rotation-scan from all three random orientations of hDOT1L (bottom). Rotation scan generated poses with minimum TCA values, from initial random orientations, that align with the DOT1L in its active state complex. b, Rotation scan TCA plots (see Extended Data Fig. 5a) for all three random starting orientations of hDOT1L in a reaction complex of nucleosome-DOT1L.
